# Competitive interactions shape brain dynamics and computation across species

**DOI:** 10.1101/2024.10.19.619194

**Authors:** Andrea I. Luppi, Yonatan Sanz Perl, Jakub Vohryzek, Pedro A. M. Mediano, Fernando E. Rosas, Filip Milisav, Laura E. Suarez, Silvia Gini, Daniel Gutierrez-Barragan, Alessandro Gozzi, Bratislav Misic, Gustavo Deco, Morten L. Kringelbach

## Abstract

Adaptive cognition relies on cooperation across anatomically distributed brain circuits. However, specialised neural systems are also in constant competition for limited processing resources. How does the brain’s network architecture enable it to balance these cooperative and competitive tendencies? Here we use computational whole-brain modelling to examine the dynamical and computational relevance of cooperative and competitive interactions in the mammalian connectome. Across human, macaque, and mouse we show that the architecture of the models that most faithfully reproduce brain activity, consistently combines modular cooperative interactions with diffuse, long-range competitive interactions. The model with competitive interactions consistently outperforms the cooperative-only model, with excellent fit to both spatial and dynamical properties of the living brain, which were not explicitly optimised but rather emerge spontaneously. Competitive interactions in the effective connectivity produce greater levels of synergistic information and local-global hierarchy, and lead to superior computational capacity when used for neuromorphic computing. Altogether, this work provides a mechanistic link between network architecture, dynamical properties, and computation in the mammalian brain.

## Introduction

A central goal of neuroscience is to understand how the architecture of the brain governs information processing. To support cognition, the brain must orchestrate the constant competition between specialised functional circuits, each arising from the cooperation of anatomically distributed regions ^1–4^. Spontaneous haemodynamics and electrodynamics provide evidence for both cooperative and antagonistic processes in the mammalian brain, exhibiting systematic and recurrent patterns of coordinated and anti-coordinated activity ^1,5–9^. Although the behavioural and physiological relevance of functional anticorrelations is well establsihed, their mechanistic origin remains unclear ^6^. How does the brain orchestrate its cooperative and antagonistic tendencies?

Interactions between brain regions unfold dynamically over a complex network of anatomical connections: the structural connectome ^10–15^. To obtain mechanistic insight about how brain structure gives rise to function, connectome-based computational models of brain activity integrate neurobiological theory and data across scales and across imaging modalities ^16–28^. Such generative models have provided growing insights about the role of regionally heterogeneous cyto-, myelo-, and chemo-architecture in shaping brain connectivity and dynamics – as well as the role of local and global network organisation of the connectome, and its variations related to evolution, development, and disease ^24,29–55^.

However, connectome-based generative models of brain activity assume that the long-range connectivity between brain regions represents cooperative interactions — that is, if region A is connected to region B, and A’s activity increases, then B’s activity will also increase. This long-standing assumption arguably arises because the main methods of reconstructing anatomical connectivity between regions (tract-tracing and diffusion tractography) produce positively-signed connectivity. Moreover, since long-range projections in the brain are excitatory, it is implicitly assumed that this representation of inter-regional interactions is appropriate (but see the recent work of Tanner and colleagues ^56^).

Yet, competitive interactions (whereby greater activity in region A leads to lower activity in B) are a ubiquitous principle of organisation in dynamical systems - whether biological, social, or artificial ^57–71^. They serve fundamental purposes such as stabilisation, feedback control, and segregating processes to avoid interference. Indeed, it is well established that the brain makes extensive use of competitive interactions at the microscale, where inhibitory synapses govern the computational and dynamical properties of neuronal circuits ^71–74^.

Here, we ask whether competitive interactions could also be present in the mammalian brain at the macroscale – and if so, how do they shape brain dynamics? We investigate this question by developing a species-specific generative model of brain dynamics that allows both cooperative and competitive interactions in its effective connectivity. This enables us to quantitatively compare models with versus without competitive interactions in terms of their faithfulness to empirical recordings of functional MRI. Does the best-fitting account of brain activity involve competitive interactions? Subsequently, we directly compare the dynamical and computational properties of cooperative-only and cooperative-plus-competitive models. Throughout, we use species-specific functional and structural data to generalise our findings about the human brain to macaque and mouse: two fundamental model organisms in translational neuroscience.

To foreshadow our results, we find that across human, macaque, and mouse the architecture of our best-fitting models consistently combines modular cooperative interactions with long-range competitive interactions, achieving up to two-fold improvement in the fit between simulated and empirical functional connectivity. The resulting dynamics are also more synergistic and hierarchical, in line with empirical observation, and exhibit more realistic values of metastability, alternating between periods of integration and segregation. Although such properties are not explicitly optimised for, they emerge spontaneously as a consequence of including competitive interactions. We conclude that our best mechanistic account of how brain structure gives rise to brain function should include competitive interactions.

## Results

We set out to investigate the presence and dynamical role of competitive interactions between regions of the mammalian brain. To obtain mechanistic insight, we use computational whole-brain models of brain activity. At their core, such generative models comprise two key ingredients: a mathematical representation of local dynamics, and a wiring diagram of inter-regional coupling ^20,28,40,75^. Specifically, our model of local dynamics consists of nonlinear Stuart-Landau oscillators poised near (but just below) the critical Hopf bifurcation ^76^. The edge-of-bifurcation dynamical working point of the Hopf model was found to produce the most faithful representation of the functional connectivity and dynamical properties of the brain ^77–80^ (Fig.1). To obtain species-specific models, we couple these oscillators according to distinct sources of data: subject-specific in-vivo diffusion tractography (human ^81^); axonal tract-tracing (mouse; ^82^); and diffusion tractography augmented with axonal tract-tracing from the CoCoMac database (macaque; ^83^). Each model is fitted to species-specific functional MRI recordings at the single-subject level (human: N=100 ^81^; macaque: N=19 ^84^; mouse: N=10 ^85^) (see Methods).

**Figure 1.**
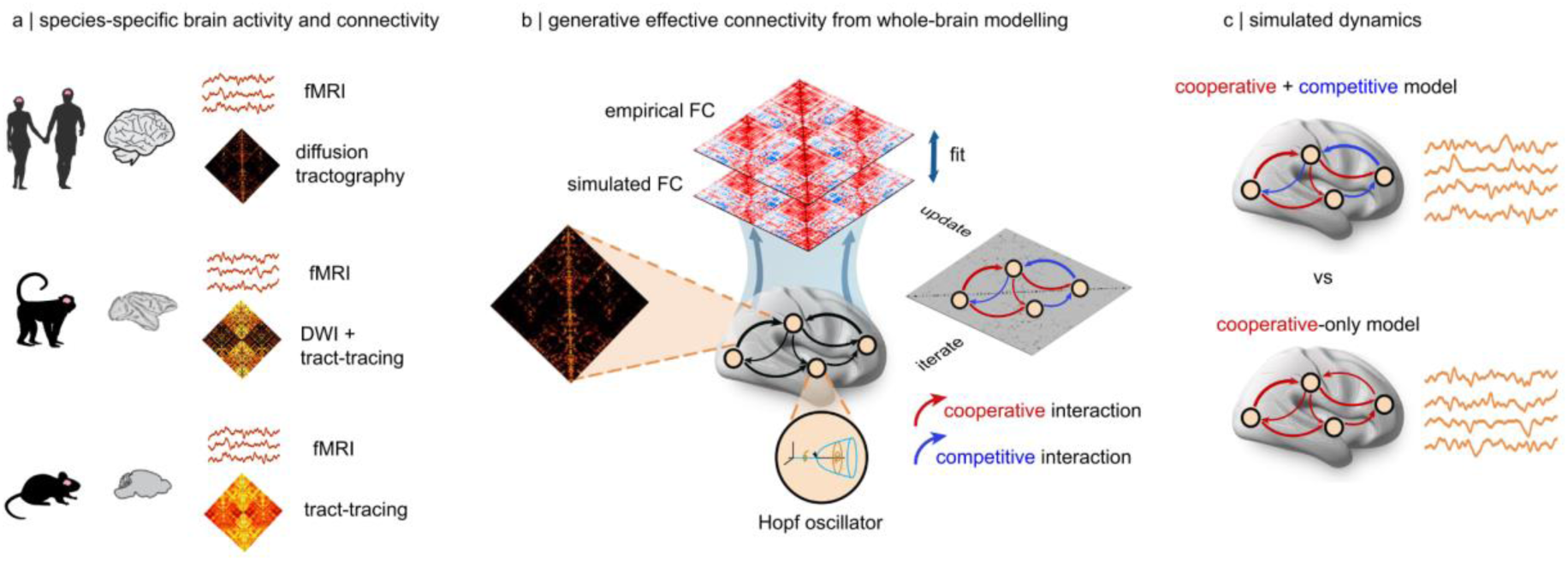
Whole-brain models generate brain activity from species-specific structural connectivity. (**a**) We develop dedicated computational models of the human, macaque, and mouse brains, based on species-specific structural connectivity: from each individual’s diffusion tractography (human); from diffusion tractography augmented with tract-tracing from the CoCoMac database (macaque); and from full tract-tracing (mouse). To facilitate comparison of results between human and other species, the structural connectomes of macaque and mouse were symmetrised, to avoid imposing structural asymmetries. Each model is fitted to species-specific functional MRI recordings at the single-subject level (human: N=100; macaque: N=19; mouse: N=10). (**b**) Overview of modelling procedure. Each brain region is modelled as a Hopf oscillator poised near (but just below) the critical bifurcation, and regions are interconnected according to the wiring diagram specified by the structural connectivity. Connection weights are then iteratively and individually updated to improve the fit between empirical and simulated functional connectivity of each individual (both with and without lag). For the model that allows competitive interactions, the sign of connections is also allowed to vary. This means that competitive (i.e., negative-signed) effective connectivity is allowed, but not imposed. After convergence, the recovered weights indicate the level of coupling between regions that most faithfully reproduces the empirical FC, thereby representing the best estimate of how SC generates FC (hence the term “generative” effective connectivity). (**c**) After convergence, the generative effective connectivity obtained from the cooperative-only and cooperative-plus-competitive models of each individual are used to generate brain activity, whose dynamical and computational properties are compared between the two models.

Crucially, in whole-brain modelling studies it is common to optimise the model by tuning regional or global free parameters (e.g. a global scaling factor for the entire SC). However, here we adopt a more recent approach that allows the connection weights themselves to vary individually ^78,86^. This method is more powerful because, rather than simply identifying the best-fitting value of a global free parameter, it identifies the level of coupling between individual regions that most faithfully reproduce the observed FC between them. In this way, this approach turns the initial structural connectivity into an “generative” effective connectivity (GEC) ^78,86^.

### Competitive interactions in the effective connectivity generate more realistic structure-function relationships

Equipped with species-specific functional MRI recordings, and species-specific structural connectomes, we begin by asking: does the most faithful account of brain activity involve competitive interactions? To allow competitive interactions in the model, we allow connections in the effective connectivity matrix to take both positive and negative values – whereas the cooperative-only model is obtained by only allowing positive values of effective connectivity.

We refer to positive-valued interactions in the generative effective connectivity as “cooperative”, because their net effect is that the more region A is active, the more its downstream neighbour B will have its activity increased, proportionally to the strength of the connection between A and B. Consequently, we refer to negative-valued interactions in the generative effective connectivity as “competitive”, because greater activity in region A leads to lower activity in B, proportionally to the strength of the negative connection between them – and vice-versa, meaning that activity in one region acts to suppress the other.

Crucially, we do not specify a priori which connections (if any) should be competitive rather than cooperative. Rather, the sign and weight are determined in a fully data-driven manner by the model. Therefore, we start by assessing whether, given the opportunity to use competitive interactions, the model will rely on them to improve its performance. Performance in this context refers to the degree to which the spatial organisation of empirical functional connectivity is reproduced. Thus, the effective connectivity produced by the model represents our best inference of how structure generates function. If any competitive interactions are observed in the effective connectivity, it means that they contribute to accounting for the relationship between structure and function. The null hypothesis is therefore that the proportion of competitive interactions will not differ significantly from zero.

Our results show that we can conclusively reject this null hypothesis. When generative effective connectivity is allowed to take negative values (i.e. the model is allowed to include competitive interactions), the model consistently takes advantage of this possibility (Figure S1 and Supplementary Tables 1-3). Across species, approximately 25-40% of edges in the generative effective connectivity are negative (Human: 25% ± 8%; Macaque: 38% ± 7%; Mouse: 28% ± 4%) (Figure S1a). Notably, we found negative-valued edges in the effective connectivity of every single individual (Figure S1a). We emphasise that this is not trivial: the model is *allowed* to have negative effective connectivity, but it is not *obligated* to do so, if doing so would decrease the fit. It is entirely plausible that the models with and without competitive interactions could have ended up producing identical-looking networks of generative effective connectivity, if it had been the case that competitive interactions serve no useful role for fitting the zero-lag and lagged FC. However, this is not what we observed empirically. Instead, the model consistently settles on having negatively valued effective connectivity, in its effort to best capture the empirical FC.

This model inversion procedure indicates that within the present modelling framework, our best-fitting account of how function arises from structure should incorporate competitive interactions. Crucially, note that the model is optimised by updating each structural connection individually, to reduce the discrepancy between the value of the corresponding entry in the simulated FC versus the empirical FC. However, each effective connection also plays a more global role, because A’s inputs to B spread to B’s neighbours in turn. Conceivably, updating the effective connection between A and B could have detrimental effects on the FC between A and a third region C that is connected to B, such that optimisation of individual structural connections could lead to a globally sub-optimal solution.

To dispel this possibility, we directly compare the outputs of two models: one that allows both cooperative and competitive interactions, and one that only allows cooperative interactions (corresponding to the traditional Hopf whole-brain model). Both models (with or without competitive interactions) are optimised to reproduce empirical FC and lagged FC, and allowed to run until no further improvement is observed. Note that allowing competitive interactions is not an additional free parameter, but rather an expansion of the range of an existing parameter of the model.

Across all three species, we find that the improvement in model fit from competitive interactions is not just local, but global. Specifically, we observe a significant increase in the model’s ability to reproduce empirical FC (quantified as the correlation coefficient between simulated and empirical FC, which is commonly used in whole-brain modelling studies): both at the group level (Fig.2a,b) and even at the level of individual subjects (Fig.2c; Human: mean (SD) = 0.42 ± 0.067 for cooperative-only; 0.70 ± 0.11 for cooperative+competitive; t(99) = −25.54; p < 0.001; Hedge’s *g* = −3.04; Macaque: mean (SD) = 0.42 ± 0.079 for cooperative-only; 0.69 ± 0.08 for cooperative+competitive; t(18) = −14.24; p < 0.001; Hedge’s *g* = −3.36; Mouse: mean (SD) = 0.73 ± 0.058 for cooperative-only; 0.87 ± 0.05 for cooperative+competitive; t(9) = −7.44; p < 0.001; Hedge’s *g* = −2.48).

**Figure 2.**
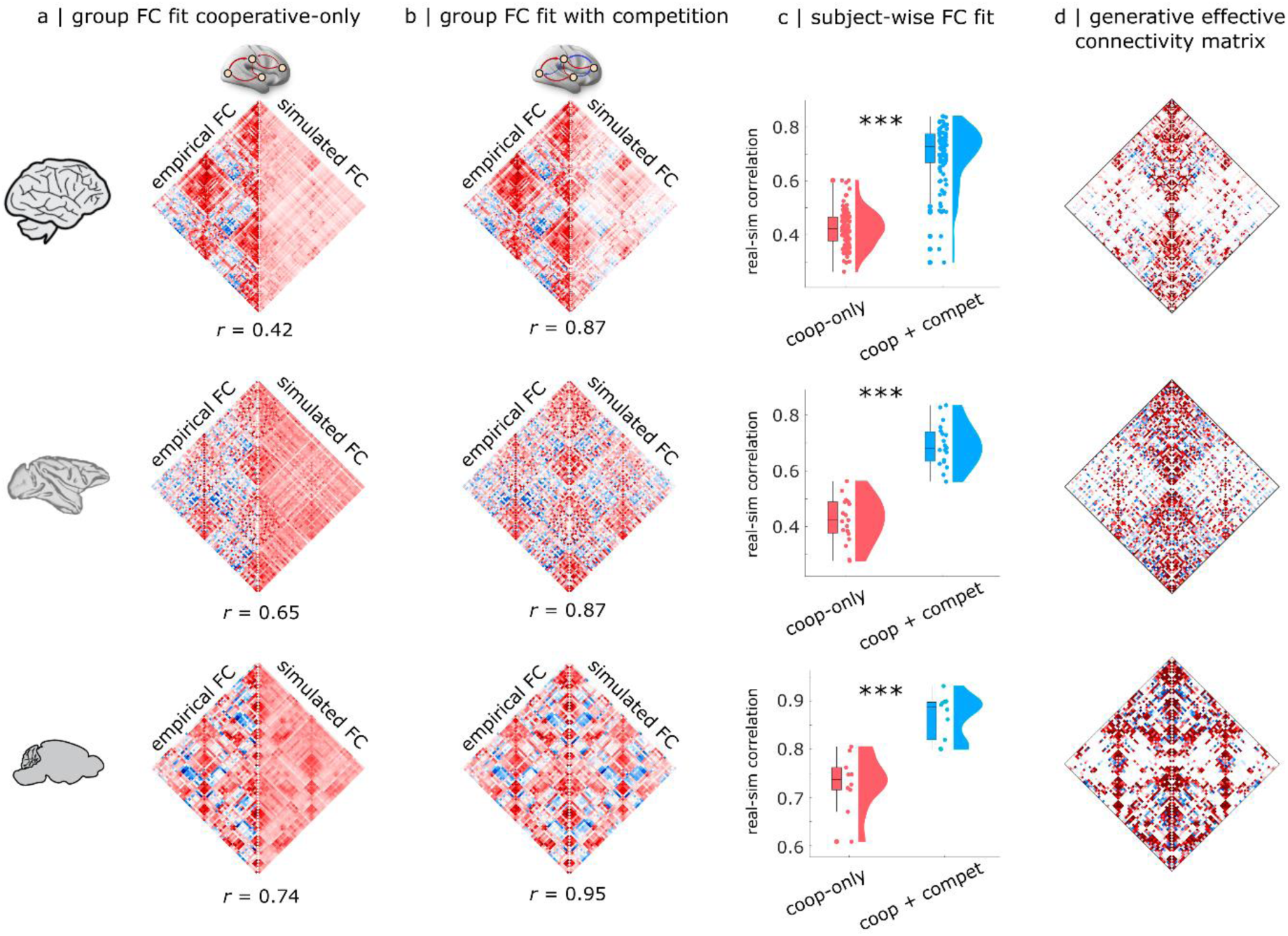
Generative competitive interactions lead to superior model fit across mammalian brains. (**a**) Species-specific matrix showing the group-average empirical (left half) and simulated (right half) functional connectivity, for the model with only positive interactions. (**b**) Species-specific matrix showing the group-average empirical (left half) and simulated (right half) functional connectivity, for the model allowing both cooperative and competitive interactions. The correlation coefficient between empirical and simulated group-wise FC is shown underneath each matrix. (**c**) Fitting quality (correlation between empirical and simulated FC) at the level of individual subjects is significantly higher when negative interactions are allowed. (**d**) Species-specific matrix of generative effective connectivity (GEC) averaged across subjects, for the model allowing negative interactions. ***, *p* < 0.001.

This improvement in global fit is so remarkable as to be visible with the naked eye: for both the mouse and macaque, the group-wise FC generated by the model with both cooperative and competitive interactions in the effective connectivity is visually indistinguishable from the empirical FC (Fig. 2b). This is confirmed numerically, with empirical and simulated group-wise FC exhibiting correlations of 0.87 (macaque) and 0.95 (mouse). In contrast, clear differences between real and simulated FC are noticeable for the model with positive-only effective connectivity (Fig. 2a). Although less visually striking, the human results are remarkable because they show the largest improvement: the model with competitive interactions more than doubles the group-level correlation between empirical and simulated FC, going from 0.42 (traditional Hopf model with cooperative-only interactions) to 0.87 (cooperative-plus-competitive). This high correlation between empirical and predicted FC compares favourably to other generative, statistical, and communication models for linking structure and function ^15^.

In turn, the presence of competitive interactions in the effective connectivity (Fig. 2d) translated to a substantially higher prevalence of negative edges (anticorrelations) in the functional connectivity, much closer to the level observed in empirical FC across species (Figure S1a,b; see Supplementary Tables 1-3 for full statistical reporting). To be clear, the model with exclusively positive effective connectivity can also produce a small number of negative functional connectivity (Figure S1b). However, the prevalence of negative FC is substantially higher (and close to empirically observed levels) in the model that also allows competitive interactions (Figure S1 and Supplementary Tables 1-3).

### Competitive interactions lead to models with greater subject-specificity

Additionally, we find that the model with competitive interactions is not only better able to reflect the empirical FC that it is intended to model, but is also better at distinguishing between the FC patterns of different individuals (Fig. 3). Arguably, a model could be very good but also very un-specific, if it exhibited equally good fit to the FC of every individual — for example, by only capturing elements that are common across all brains, while disregarding subject-specific ones. This concern may be especially prominent for our macaque and mouse models, which are based on species-specific connectomes rather than individual-specific ones (whereas the human models were fitted using individualised structural connectomes).

**Figure 3.**
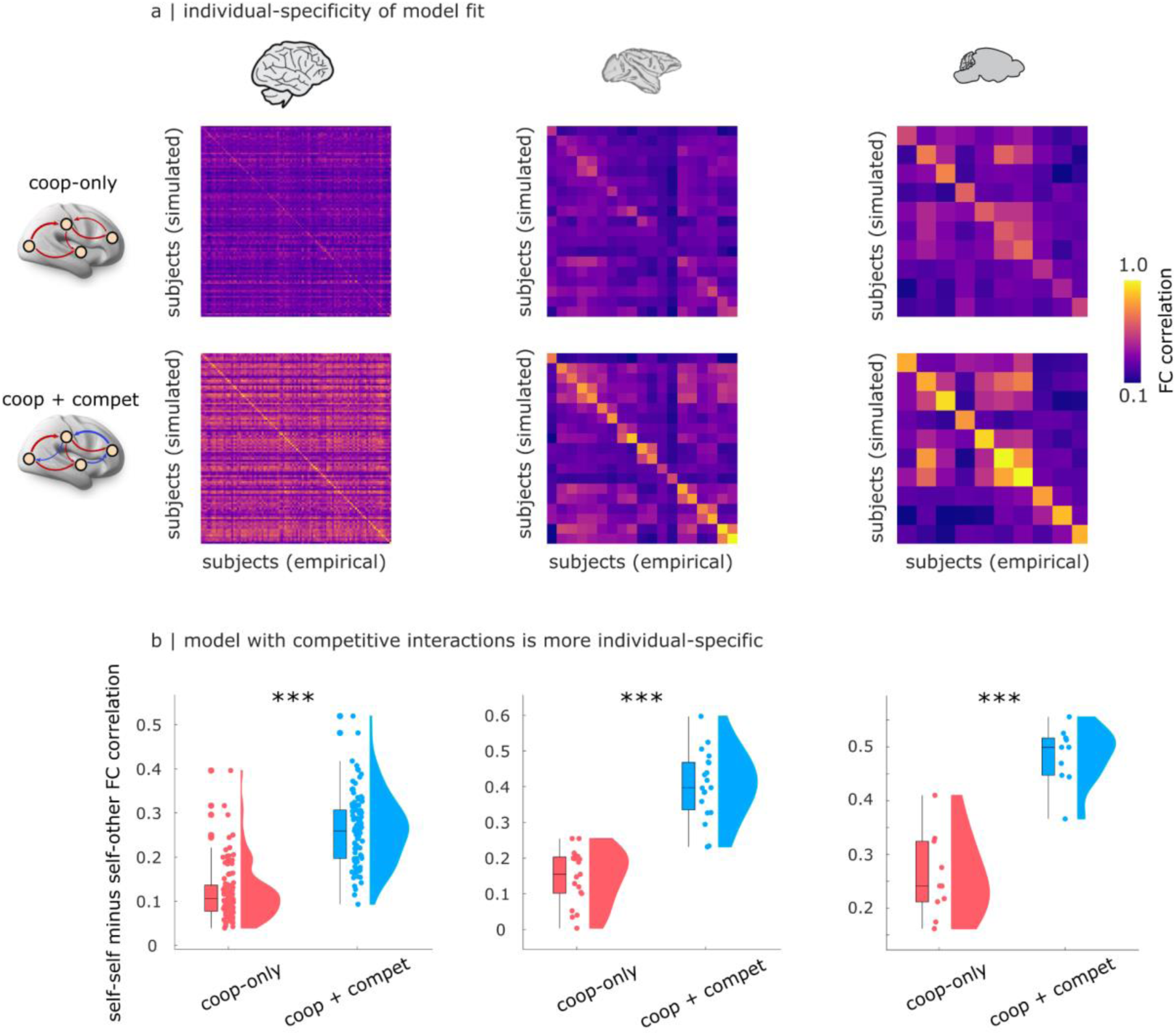
Competitive interactions produce models with greater subject-specificity. (**a**) Matrix of FC similarity (correlation) between each empirical subject (columns) and each simulated subject (rows). Brighter diagonal therefore reflects greater success of the commonly used model-fitting criterion of similarity between a subject’s own empirical and simulated FC (as reported in Fig.2). (**b**) Differential identifiability is defined as the difference between self-self similarity of FC (diagonal elements in panel (a)) and mean self-other similarity of FC (off-diagonal elements), such that greater difference implies more individual-specificity. In other words, it quantifies the cost of mis-matching individuals. When this cost is low, there is low subject-specificity. In the “brain fingerprinting” literature, differential identifiability quantifies the ease of telling apart different individuals based on their FC; here we apply it to quantify the individual-specificity of brain models, such that greater differential identifiability indicates that there is a larger drop in similarity between simulated and empirical FC when a model is matched to the wrong individual. We find that models with both cooperative and competitive interactions are significantly more individual-specific. ***, *p* < 0.001.

However, we show that this is not the case. We quantify this phenomenon using the measure of “differential identifiability” from the literature on brain fingerprinting, which is defined as the mean difference between self-self and self-other similarity of FC ^87^. In other words, this measure quantifies the cost of mis-matching individuals in terms of loss of model fit. For both models (with and without competitive interactions), we find that differential identifiability is always positive (Fig. 3a): the model reproducing an individual’s FC exhibits greater affinity for the empirical FC of that individual, than for the FC of other individuals. Crucially, however, we also find that the model with competitive interactions exhibits significantly larger differences between self-self and self-other FC fit (Fig.3b; Human: mean (SD) = 0.12 ± 0.06 for cooperative-only; 0.26 ± 0.07 for cooperative+competitive; t(99) = −16.97; p < 0.001; Hedge’s *g* = −2.00; Macaque: mean (SD) = 0.15 ± 0.075 for cooperative-only; 0.40 ± 0.09 for cooperative+competitive; t(18) = −15.42; p < 0.001; Hedge’s *g* = −2.88; Mouse: mean (SD) = 0.25 ± 0.077 for cooperative-only; 0.48 ± 0.05 for cooperative+competitive; t(9) = −7.01; p < 0.001; Hedge’s *g* = −3.18). In other words, there is a larger drop in similarity between simulated and empirical FC when a model is matched to the wrong individual. This effect is consistently observed across species. Note that for the human data, both the cooperative-only and the cooperative-plus-competitive model are based on individual SC: therefore, the use of individual SC cannot be driving differences in the two model’s performance. Thus, we conclude that the model with competitive interactions is significantly more individual-specific.

Do positive and negative edges in the effective connectivity differ only in terms of sign, or are there systematic differences in their organisation? Across species, competitive connections can be observed even in the group-average effective connectivity (Fig.2d,e), indicating the consistency of their placement across individuals. Additionally, we find that negative effective connections are less modular than positive connections (Figure S2b; Supplementary Tables 1-3), and exhibit a lower clustering coefficient (Figure S2a; Supplementary Tables 1-3) — in other words, it is less likely that negatively-interacting neighbours of a node will themselves be negatively-interacting. Negative connections are also found to be weaker and longer than positive ones (Figure S2c-d; Supplementary Tables 1-3). Overall, the picture emerges of a modular network of strong positive ties, alongside long-range, diffuse negative ties.

### Dynamical consequences of competitive interactions in the effective connectivity

We have shown that competitive interactions in the effective connectivity shape the *spatial* organisation of functional connectivity. However, mammalian brains also exhibit rich dynamics, giving rise to prominent patterns of *temporal* signal coordination ^88–93^. Therefore, the question arises: do competitive interactions also shape brain dynamics?

Remarkably, we find that in addition to achieving substantially better spatial similarity with the empirical FC, the model with competitive interactions also exhibits more realistic dynamics, which it was not explicitly optimised for. To demonstrate this, we use the effective connectivity recovered by each model (with and without competitive interactions) to simulate regional brain activity (hence switching from inverse modelling to forward modelling ^21^). One aspect of brain dynamics that is often studied is metastability, which at the macroscale is typically operationalised as the temporal standard deviation of the Kuramoto order parameter (KOP ^77,94^; see ^95^ for an extensive discussion of metastability and its alternative signatures in the literature). Since the KOP quantifies instantaneous synchrony, its variability over time quantifies the tendency of the brain to alternate between periods of synchronisation and desynchronisation, reflecting the co-existence of integrative and segregative tendencies ^95^. Results show that metastability (*std(KOP)*) exhibits unrealistically high values for the model with positive-only interactions in the effective connectivity, in each of the three mammalian species considered (Fig.4a). In contrast, including competitive interactions lead to a more balanced level of metastability, closer to empirical values (Fig.4a; see Supplementary Tables 1-3 for full statistical reporting).

**Figure 4.**
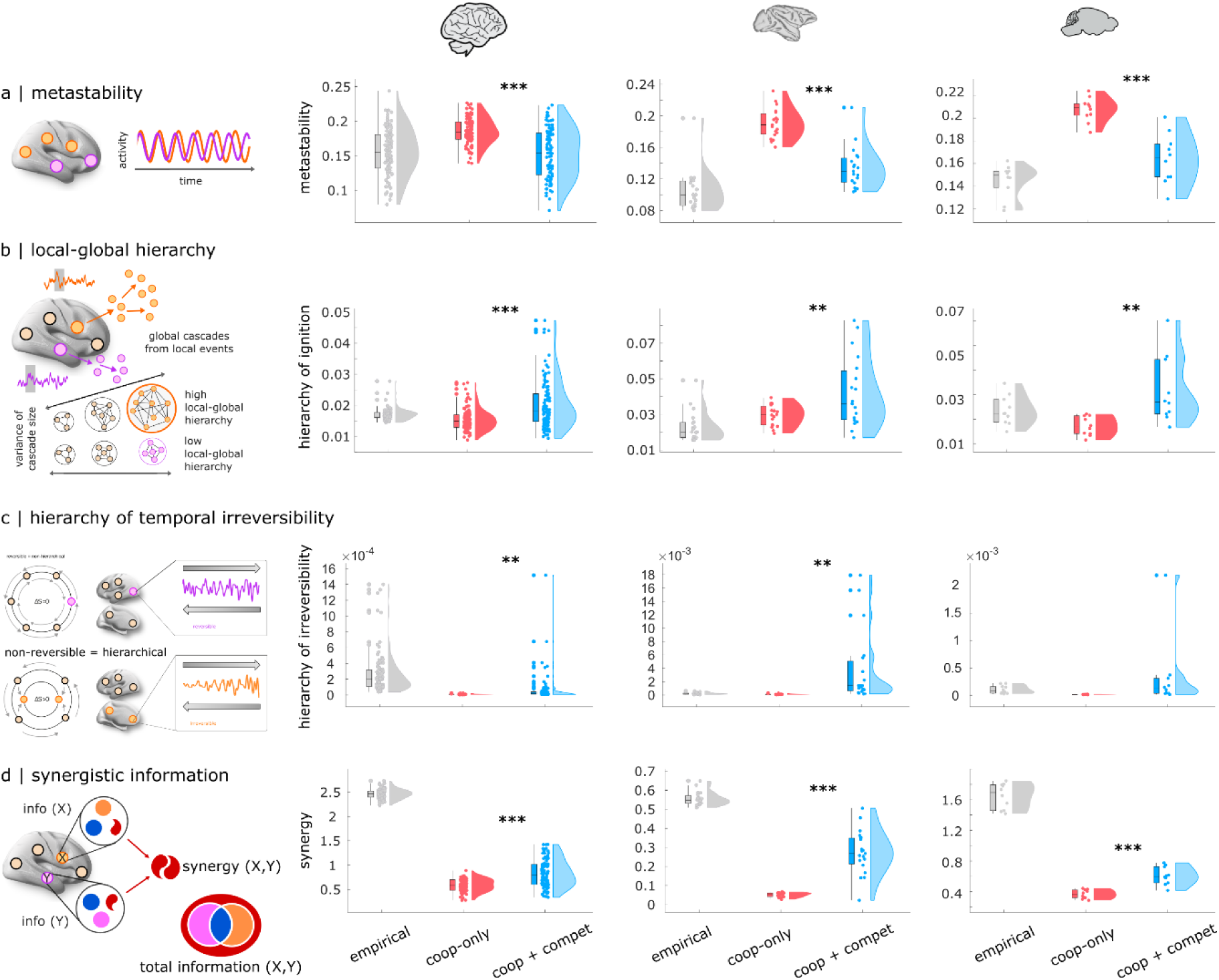
Dynamical consequences of competitive interactions in the effective connectivity. (**a**) **Metastability.** Metastability (here operationalised as the temporal standard deviation of the Kuramoto order parameter) reflects the temporal alternation of synchronisation and desynchronisation across the brain. (**b**) **Local-global hierarchy.** Intrinsic-driven ignition (IDI) is obtained by identifying “driver events” (unusually high fMRI spontaneous activity) and measuring the number of co-occurring events. A measure of local-global hierarchy is obtained by calculating the variability across regions of their average IDI size over time, such that the brain is more hierarchical when there is greater disparity across the size of elicited intrinsic neural events. (**c**) **Hierarchy of temporal irreversibility.** Level of hierarchy is given by the level of asymmetry of interactions between brain regions arising from the breaking of the detailed balance. A non-hierarchical system is depicted, being in full detailed balance and thus fully reversible over time. In contrast, asymmetry of the interactions results in a hierarchical organisation. (**d**). **Synergistic information.** The total information jointly carried by two variables X and Y (e.g., two brain regions) can be exhaustively decomposed into information that is redundantly carried by both variables (blue); or uniquely by each (yellow and orange); or synergistically by considering the two variables together (red). Each column displays results for a different species. *, *p* < 0.05; **, *p* < 0.01; ***, *p* < 0.001. Full statistical reporting for the comparisons between cooperative-only and cooperative+competitive models is provided in Supplementary Tables 1-3.

Our results also show that the model with competitive interactions in its generative effective connectivity exhibits a more hierarchical organisation, operationalised in two different ways. “Intrinsic-driven ignition” refers to the capacity of local events of unusually high activity to propagate globally throughout the brain, which is thought to be necessary for allowing integration and differentiation to co-exist in the brain ^45,96,97^. Building on this principle, a measure of local-global hierarchy can then be obtained by comparing the variability of regions’ capacity to ignite global activity: the brain is more hierarchical when there is greater disparity across regions for the size of elicited intrinsic neural events. In other words, local-global hierarchy is operationalised in terms of covering a broader range from local to global size of intrinsically-originated events (Fig.4b): when event sizes are all near-equal (whether all very localised, or all very global), there is low hierarchy, whereas when there are wide differences in event size across regions, then there is high level of local-global hierarchy in the brain ^45,96,97^. We find that this measure is consistently increased for the model with versus without competitive interactions (Fig.4b; Supplementary Tables 1-3).

Another way of conceptualising hierarchical organisation in the brain is related to the temporal (ir)reversibility of brain activity, which gives rise to asymmetric interactions between brain regions ^78,98^. When interactions are symmetric then sending and receiving of information are in equilibrium for each region, while when interactions are asymmetrical then directionality exists, such that regions will differ in terms of whether they have a preference for sending or receiving signals from other regions. Hence, the greater the variability in this send-receive asymmetry across regions, the more the system’s organisation is hierarchical — i.e. departing from the equilibrium of a flat organisation ^78^. Our results show that the presence of competitive interactions in the generative effective connectivity induces more hierarchical organisation of asymmetry, with greater divergence in regions’ preference for sending or receiving information, being closer to what is observed in empirical brain dynamics (Fig.4c). We note that both the models with and without competitive interactions include lag-1 FC in their fitting function, thereby incorporating some aspect of directionality and irreversibility, as previously recommended ^78^. Nonetheless, we see that the model with competitive interactions is better capable of reflecting variability of send-receive asymmetry across regions, compared with the model that only allows positive values of effective connectivity (Fig.4c; see Supplementary Tables 1-3 for full statistical reporting).

Finally, we consider the prevalence of synergistic information in the intrinsic dynamics of the brain ^99^. In a dynamical system such as the brain, activity is not random: even at rest (i.e., in the absence of any explicit task), the future state of the brain’s spontaneous activity will depend on its past state. This means that the past trajectory of a region’s activity holds information about its future state, known as “time-delayed” information ^100^. However, brain regions are interconnected and continuously interacting, such that the activity of region X may also influence the future activity of region Y, and vice-versa. Mathematically, the total information jointly carried by two variables X and Y (here, two brain regions) can be exhaustively decomposed into information that is carried redundantly by both variables (such that it is equally available from each of them); information that is carried uniquely by only one variable; and finally, information that is carried by the two variables synergistically ^99,101–106^. Synergistic information is available only when both variables are considered jointly, but not when considering either variable in isolation - thereby reflecting the extent to which interactions in the system are “greater than the sum of their parts”. It was recently shown that synergy is associated with higher order cognitive operations and evolutionary expansion of the human brain, being also diminished when consciousness is lost due to pharmacological or pathological conditions ^99,101,107^. Our results show that although both models fall short of the level of synergy observed in empirical brains, the model with competitive interactions produces significantly higher values of synergy in its dynamics, consistently exceeding the model with positive-only interactions, across all three mammalian species considered (Figure 4d and Supplementary Tables 1-3).

### Competitive interactions increase the match between simulated brain activity and canonical cognitive operations of the human brain

Up to this point, we have found that allowing the generative effective connectivity to include competitive (negative-valued) interactions leads to significantly greater fidelity to the spatial and temporal organisation of the mammalian brain. Ultimately, the goal of spatial and temporal coordination between brain regions is to support cognition. To support different cognitive roles, brain regions co-activate together to form specialised functional circuits. Such coherent patterns of co-activation can be observed even in the spontaneous activity of the brain, suggesting that they are an intrinsic feature of brain functional organisation ^108,109^. Does our model also exhibit the spontaneous emergence of coherent functional circuits?

To quantify how well the instantaneous patterns of brain activity match the brain patterns associated with canonical cognitive operations, we can use our recently introduced “cognitive matching” procedure ^110^. The cognitive matching score is computed as the best spatial correlation between brain activity and 123 brain maps obtained by meta-analytic aggregation of thousands of human neuroimaging studies from the NeuroSynth engine and Cognitive Atlas (Fig.5a) ^110–115^. For each individual (real or simulated), an overall index of the quality of cognitive matching is obtained by averaging the cognitive matching scores across the entire scan duration. Higher match to NeuroSynth meta-analytic maps indicates greater alignment of spontaneous brain activity with brain maps from the cognitive neuroimaging literature. In somewhat poetic terms, cognitive matching quantifies how “mind-like” the observed brain patterns are. Indeed, we have recently shown that the quality of cognitive matching declines sharply in response to anaesthesia, when consciousness and cognition are suppressed ^110^.

**Figure 5.**
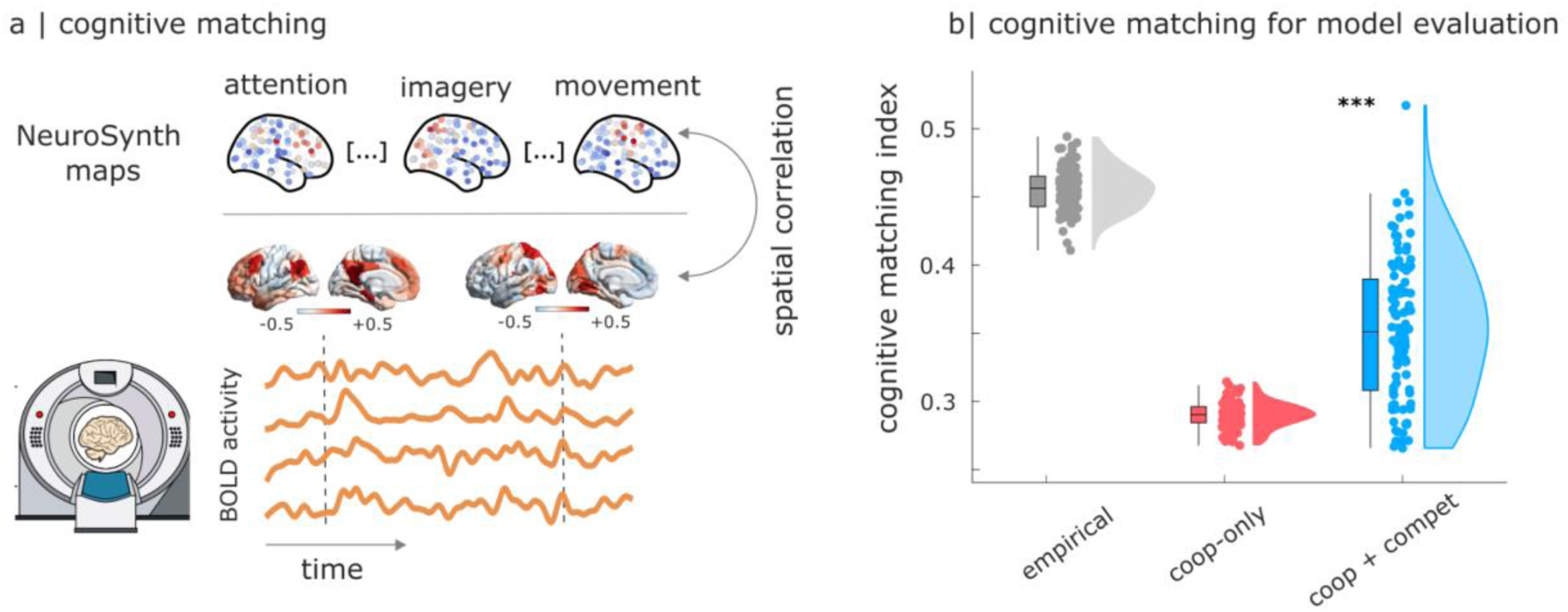
Competitive interactions increase the match between simulated brain activity and canonical cognitive operations of the human brain. (**a**) At each point in time, the cognitive matching score is computed as the best spatial correlation between brain activity and 123 NeuroSynth meta-analytic brain maps. For each individual (real or simulated), an overall index of the quality of cognitive matching is then obtained by averaging the cognitive matching scores across the entire scan duration. Higher NeuroSynth matching indicates greater alignment of spontaneous brain activity with brain maps from the cognitive neuroimaging literature. (**b**) Model with negative effective connectivity produces patterns of brain activity that are significantly more realistic than when only positive effective connectivity is allowed. ***, *p* < 0.001.

Thus, through cognitive matching we can assess whether our computational models co-activate regions that we know belong to the same cognitive circuit in the real brain (as indicated by meta-analytic co-activation). Our results show that the cognitive matching index of the model with competitive interactions is significantly higher than the one of the model with only cooperative interactions (mean (SD) = 0.29 ± 0.01 for cooperative-only; 0.35 ± 0.05 for cooperative+competitive; t(99) = −13.96; p < 0.001; Hedge’s *g* = −1.67), eliciting a substantial shift towards the levels of cognitive matching observed for the real human brain (Fig.5b). In other words, the model with competitive interactions is more capable to co-activate regions that belong to the same functional circuit, as defined in a data-driven manner from thousands of cognitive neuroscience experiments. (Note that, at present, NeuroSynth is only available for human neuroimaging studies, making this type assessment of model quality only possible for the case of human brain activity).

### Competitive interactions increase computational capacity

Despite being termed “functional”, the functional connectivity need not reflect any genuine function, in the sense of ongoing cognition ^116^. Our cognitive matching procedure improves on this shortcoming, by associating specific patterns of brain activity with specific cognitive operations. An alternative, complementary strategy is the recently introduced approach of connectome-based reservoir computing ^117–119^. Under this framework, a connectome becomes the network that is used for a neuromorphic architecture performing a specific task, such as memorising some time-series. Notably, the task is performed *in silico* by the connectome, rather than in vivo by a person or animal in the scanner. Task performance can then be used as a read-out for the computational capacity of the network (here, connectome), reflecting its suitability for performing useful computation. Here, following the workflow established by ^118,119^ we use the reservoir to perform a memory task, with each individual’s generative effective connectivity as the network. For each species, input nodes are defined as the visual regions of the cortex, and output nodes are defined as the motor regions, reflecting their biological roles (Fig.6a).

**Figure 6.**
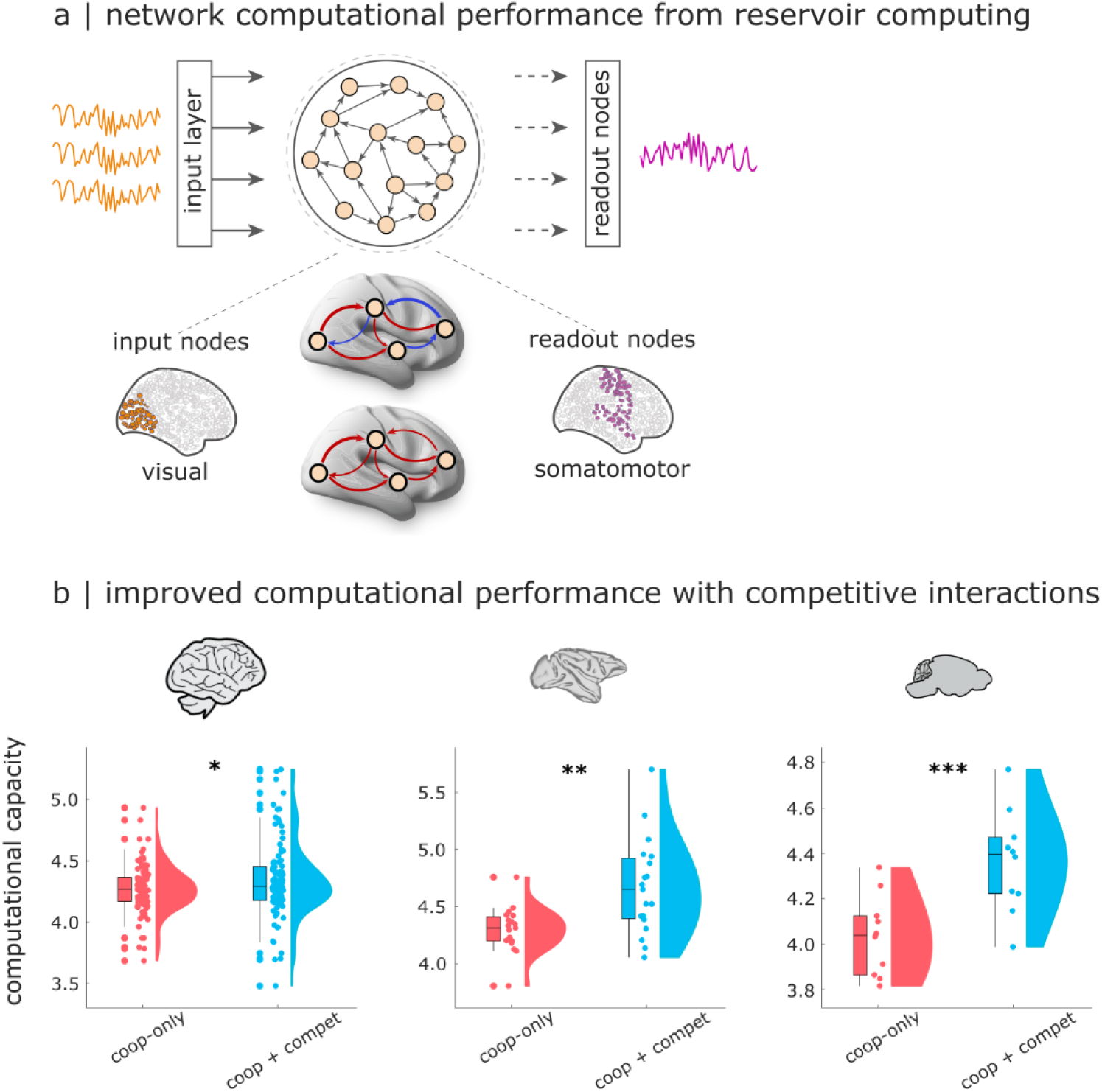
Superior computational performance of connectome-based neuromorphic networks with competitive generative interactions. (**a**) The reservoir computing architecture consists of an input layer, followed by a *reservoir* (nonlinear recurrent neural network), and a linear readout module. Here, we endow reservoirs with effective brain connectivity patterns and evaluate the effect of competitive generative interactions on computational performance using a memory task. In this task, the readout module is trained to reproduce a time-delayed version of a random input signal, measuring the reservoir’s ability to encode past stimuli. We choose visual cortical regions as input nodes and somatomotor regions as output nodes to reflect their respective functional roles. (**b**) Memory capacity of the connectome-based reservoir is significantly higher in the presence of negative effective connectivity, across all three mammalian species considered. *, *p* < 0.05; **, *p* < 0.01; ***, *p* < 0.001.

Remarkably, our results reveal that the effective connectivity networks obtained from a model with competitive interactions exhibit a superior computational capacity, consistently outperforming the networks obtained with cooperative-only interactions (Fig.6b; Human: mean (SD) = 4.29 ± 0.19 for cooperative-only; 4.36 ± 0.30 for cooperative+competitive; t(99) = −2.22; p = 0.029; Hedge’s *g* = −0.28; Macaque: mean (SD) = 4.27 ± 0.27 for cooperative-only; 4.63 ± 0.42 for cooperative+competitive; t(18) = −3.17; p = 0.005; Hedge’s *g* = −0.97; Mouse: mean (SD) = 4.03 ± 0.17 for cooperative-only; 4.36 ± 0.02 for cooperative+competitive; t(9) = −7.60; p < 0.001; Hedge’s *g* = −1.53). Once again, this effect is observed consistently across species. In other words, in addition to better matching the functional patterns of the human brain, the model with competitive interactions can also be used to execute an actual task with greater performance.

### Robustness analysis

Throughout this work, we have consistently replicated our results in three independent datasets pertaining to three distinct mammalian species, demonstrating their generality. In this section, we summarise the results of several additional control analyses that we performed to ensure the robustness of our findings.

First, it is important to remark that the presence of anticorrelations in our three empirical datasets is not due to the preprocessing step of global signal regression (GSR). Indeed, since it is known that this procedure can introduce artificially high levels of anticorrelations in the FC ^6,120^ we elected not to use GSR for any of our datasets. Therefore, the presence of empirical anticorrelations is unrelated to GSR. Nonetheless, in Figure S3 we show that our main results are successfully replicated when GSR is used, achieving an even better fit between real and simulated human FC.

We also confirm that the human results can be replicated using a different, more fine-grained parcellation of the cortex (Schaefer-200 ^121^), and also with addition of subcortical structures from the Tian atlas ^122^ (Figure S4). Since diffusion tractography is known to have difficulty resolving interhemispheric connections between homotopic regions ^123–126^, we also replicate our human results after adding homotopic connections - which does result in a small improvement (Fig. S5). Similarly, although we symmetrised the macaque and mouse connectomes to facilitate comparison with the human data, we verify that the macaque and mouse results can be replicated if the asymmetries (and for mouse, the fully-dense nature) of the structural connectome are preserved (Figures S6-S7).

Additionally, we investigate if the results could be driven by a difference in overall connectivity between the resulting cooperative-only and cooperative-plus-competitive models. For this, we consider a different way of comparing these models: rather than fitting a cooperative-only model in the traditional way (whereby updates that would make a connection negative are disallowed), we instead obtain the effective connectivity from the cooperative-plus-competitive model, and then simply take the absolute value such that all competitive interactions are now cooperative. As a result, the magnitude of inter-regional coupling between each pair of regions is identical for the cooperative-only and cooperative-plus-competitive models, which was not guaranteed when fitting the models separately. Nevertheless, results across all three species show that if competitive interactions are turned cooperative, the resulting FC loses structure and performance declines dramatically, accompanied by a loss of dynamical realism (Figures S8-S10). Note that this result should not be surprising: this version of a cooperative-only model *could* have been produced by the model optimisation, but was not, because it did not lead to good fit. This analysis confirms that the competitive nature of the interactions is important, not merely their number and weight.

Finally, we also show that an alternative measure of global FC fit, the structural similarity index (SSIM) that is commonly used for comparing images ^127^, is also significantly improved for the model with both cooperative and competitive interactions in the effective connectivity (Figure S11 and Supplementary Tables 1-3).

## Discussion

Altogether, we examined the dynamical and computational relevance of competitive interactions in the mammalian connectome. We used generative computational modelling to integrate multi-modal data about brain structure and function across human, macaque, and mouse brains. Consistently across species, our best mechanistic account of how brain network structure gives rise to function combines modular cooperative interactions with diffuse, long-range competitive interactions. Models with competitive interactions are more individual-specific, and achieve excellent fidelity to the spatial coordination and temporal dynamics of the empirical brain, including properties that were not explicitly optimised but rather emerged spontaneously. Below we discuss the implications of these findings while outlining future work.

### Competitive interactions enable superior realism across space and time

Our species-specific computational models revealed that allowing the generative effective connectivity to include competitive as well as both cooperative interactions produces consistently superior fit to the empirical functional connectivity, both at the group level (with up to 0.95 correlation between real and simulated FC) and at the level of individual subjects. Remarkably, upon introducing competitive interactions the increase in the model’s explicit fitting objective (spatial organisation of functional connectivity) was accompanied by increased fit to several additional, *un-optimised* properties, which are dynamical rather than spatial: metastability, synergy, and hierarchical organisation. We found that our signature of metastability in the model with cooperative-only interactions is consistently higher than in real brains (consistent with similar observations from the recent voxelwise model of ^44^). Adding competitive interactions brings it closer to empirically-observed levels. A possible interpretation of this observation is that metastability (here operationalised as the temporal variability of the Kuramoto order parameter, as is common in the fMRI literature ^77,94,95^) is intended to quantify a balance between integrative and segregative tendencies. Temporal variability of the brain’s level of synchrony (KOP) indicates that high and low synchronisation co-exist and alternate over time. If only cooperative interactions are present, this can lead to periods of excessive synchrony due to regions reciprocally increasing each other’s activity level in a vicious cycle (we confirm this in Figure S12, showing that the maximum observed synchrony is significantly higher for the model without competitive interactions, often approaching total synchronisation). Competitive interactions may play a stabilising role, because as one region’s activity grows, activity of its competitively-connected neighbours will diminish, preventing runaway global activity.

For similar reasons, competitive interactions may enhance the level of local-global hierarchy by increasing the divergence between different regions’ ability to ignite global activity. Even when connections are exclusively cooperative, heterogeneity of connection strength can promote ignition-like activity ^42^. In the presence of competitive interactions, the global propagation of local events depends on the balance between cooperative interactions (which favour spreading) and competitive interactions (which have the opposite effect, suppressing the activity of downstream neighbours), meaning that regions can now vary in their overall ignition capacity alongside two dimensions, rather than just the positive one. This observation is consistent with the work of ^128^, who showed that in a system of coupled oscillators, a mixture of both positive and negative couplings is optimal for signal amplification.

Finally, synergy reflects the presence of complementary information in different brain regions’ activity, such that there is greater ability to predict the joint future state of two brain regions, when both regions are considered together rather than in isolation ^99,101^. In practice, synergy is low both when two regions are fully independent (such that neither of them has information about the other, but only about itself) but also when two regions are strongly dependent, such that the same information is present in each, without need to consider the other ^99,101^. Thus, synergy reflects a balance between dependency and independence in the system, complementing metastability (which in its operationalisation as *std(KOP)* only considers instantaneous relationships). A system with synergy is a system whose elements act as a whole rather than as disjoint parts. We found that by countering the tendency to extreme synchronisation, competitive interactions allow greater diversity of activity to be present and available for integration, manifesting as greater synergy.

Altogether, competitive interactions increase synergy and hierarchical functional organisation while exerting a moderating effect on metastability, bringing the simulated dynamics more in line with empirical dynamics of the mammalian brain. Remarkably, this increased fidelity of the model’s dynamics is accompanied by increased realism of the specific patterns of model-generated brain activity, as revealed by our newly introduced “cognitive matching” criterion. The cognitive matching index reflects the similarity between spontaneous brain activity, and brain patterns associated with canonical cognitive operations, as synthesised in a data-driven way from over 14,000 published neuroimaging studies ^110^. When applied to recordings from real human brains, cognitive matching may be understood as recovering echoes of cognition in the spontaneous activity of the brain. When translated into a criterion of model realism, as introduced in the present report, cognitive matching measures whether the model co-activates regions in a manner consistent with their membership of the same cognitive circuits. Thus, the present results suggest that competitive interactions are crucial for enabling co-activation of regions that belong to the same cognitive macro-circuit, because we are significantly less likely to observe appropriate co-activations in the absence of competitive interactions.

### Mechanistic insights about cooperative-competitive architecture of the mammalian connectome

A key advantage of our model-based approach is that we can make claims about the causal role of the competitive interactions, because we work with effective connectivity obtained from a generative model. In other words, unlike studies of functional connectivity, here we can assign a definite mechanistic meaning to the negative sign of the competitive effective interactions: they represent suppression of the target’s activity by the source, justifying the term “competitive”.

Altogether, our generative model consistently indicates that competitive interactions play a pivotal role in explaining the emergence of functional interactions from the brain’s structural connectivity, achieving unprecedented realism. This meaning of “functional” goes beyond regional co-fluctuations (what is commonly referred to as functional connectivity), instead also encompassing co-activation of brain regions in accordance with membership of the same cognitive macro-circuit, as determined by meta-analytic synthesis with NeuroSynth. The same competitive interactions also endow brain dynamics with greater synergy and hierarchy, and overall greater fidelity to the empirical dynamics. Note that only FC and lagged-FC are explicitly optimised by the algorithm, yet the remaining criteria are also significantly improved – suggesting that the model is not overfitting, but rather genuinely moving towards greater fidelity across both spatial and dynamical dimensions.

Systematic patterns can also be identified in the organisation of competitive interactions, which are consistent across species. Whereas cooperative (positive-valued) interactions in the effective connectivity tend to be strong, modular, and relatively short-range, competitive interactions are weaker but more long-range, more diffuse, and less clustered: it is less likely that competitively-interacting neighbours of a node will themselves be interacting competitively. Corroborating these results, lower clustering for negatively-weighted than positively-weighted connections was also recently reported by Tanner and colleagues ^56^.

The phenomenon of strong modular connections complemented by diffuse weaker ones is reminiscent of the “strength of weak ties”, observed in social and neuronal networks alike ^129,130^. Our present results suggest that in the mammalian brain, such weak ties may be of a predominantly competitive nature. The resulting architecture balances local cooperation (within specialised modules), and global competition - reminiscent of the “global workspace” architecture ^2,4^. We have shown that such a cooperative-competitive architecture achieves a greater range of local-to-global responses, and greater synergy in its dynamics. In the real human brain, synergy is especially prevalent in integrative regions of association cortex that support higher-order cognitive functions ^101,131,132^. Convergent evidence for an association between synergy and cognitive capacity was recently provided, showing that synergy supports flexible and efficient learning in artificial neural networks ^133^. This evidence is further corroborated by our own present results: across humans, macaques, and mice, we found that connectome-based neuromorphic networks reach the highest computational capacity when both cooperative and competitive interactions are present. It is intriguing that in addition to matching spatio-temporal properties of the real brain, competitive interactions also endow the effective connectivity with greater computational capacity – given that one of the brain’s chief functions is arguably that of enabling computation.

### Limitations and Future Directions

A major strength of the present study is the consistent replication of results across three independent datasets pertaining to three different mammalian species. However, our species-specific datasets also inevitably come with numerous differences. For the functional MRI data, differences include scanner and acquisition parameters. For the structural connectome data, the mouse SC was entirely obtained from axonal tract-tracing, which is considered the current gold-standard ^134^, whereas the macaque connectome was reconstructed by augmenting diffusion MRI tractography with existing tract-tracing data, and the human structural connectomes were reconstructed entirely from diffusion MRI tractography, as the only technique that can be applied *in vivo*. However, tractography-derived connectomes can exhibit both false positives and false negatives ^123–126^. The superiority of tract-tracing may account for the higher quality of fitting observed in mice than in humans and macaques (indeed, the mouse tract-tracing connectome – and resulting effective connectivity – display clear inter-hemispheric connections, which are notoriously difficult to resolve with diffusion tractography). On the other hand, a single macaque and mouse connectome were used for all animals, whereas for the human data we were able to use subject-specific structural connectomes, demonstrating that our modelling framework is fully applicable to the single-subject case.

Overall, the differences between datasets could represent a limitation, if the present study were focused on inter-species differences. However, our interest is rather in the consistencies across these species, which demonstrate the replicability and generalisability of our results. In this context, differences in how the various datasets were acquired may even be seen as an asset: they are evidence that our discoveries do not critically depend on the specific sampling rate or MRI parameters used, nor on the specific technique that is used to reconstruct the structural connectome, but rather reflect more fundamental properties of mammalian brain organisation.

The fact that we work with generative effective connectivity enables us to draw further inferences about the causal role of competitive interactions. However, our present modelling framework does not allow us to draw conclusions about the specific underlying neurobiology - be it direct long-range inhibition (as assumed by Dynamic Causal Modelling, which does not consider structural connectivity ^135,136^), or long-range excitation of a local inhibitory circuit, as suggested by the recent model of ^137^, or other more complex phenomena such as conduction delays or the relative contributions of cerebral blood volume and flow to the haemodynamic signal, among others ^6,138–141^. More biologically detailed models that explicitly incorporate such alternative circuits will be necessary, to distinguish between these options - possibly in combination with experimental work and approaches such as MR spectroscopy ^138,140^. Nonetheless, our observation of a role of competitive interactions in explaining macroscale brain activity (whatever their ultimate biological origin) is corroborated by two recent reports, which used different strategies. Ruffini and colleagues ^142^ fitted Ising models to binarised fMRI data, recovering a signed approximation to the structural connectome that included negative connections. Tanner and colleagues ^56^ used a multi-linear regression framework to predict the future of a region’s activity from the weighted histories of its structurally-connected neighbours, allowing them to infer the sign, weight, and direction of structural interactions. When subsequently used as the wiring diagram for a generative neural mass model, the signed SC of ^56^ outperformed the original SC at reproducing the empirical FC. Overall, both approaches converged with our own, in suggesting that our best statistical and generative accounts of structure-function relationships in the brain should incorporate negatively-signed structural connections. Identifying their neurobiological origins remains an exciting challenge for computational and systems neuroscience.

### Conclusion and Outlook

Altogether, the present work provides a mechanistic link between network architecture, dynamical properties, and computation in the mammalian brain, expanding our neurobiological understanding of the mechanisms linking structure and function. At the same time, this work represents a major advance in the development of *in silico* brain models – a topic of intense research ^54^. Generative models such as ours are particularly desirable from a clinical perspective, to develop *virtual screening tools* for potential interventions ^54^. Compared with state-of-the-art cooperative-only models, our cooperative-competitive model achieves markedly superior fidelity across diverse dimensions of brain connectivity and dynamics. This is not only observed at the group level: our models are fitted to individual human brains using their own structural and functional data. Crucially, in addition to providing a superior match to the empirical brain of each individual, our new model exhibits also greater subject-specificity. In other words, we do not simply have a better model of human brain function: we have a better model of the brain of each individual. This represents a critical advance towards the development of personalised computational models of individual patients’ brains, based on their own data. Notably, the success of our modelling framework is not limited to the human brain but rather it extends to mouse and macaque, highlighting the similarities between humans and two fundamental model organisms in neuroscience. Invasive therapeutic interventions are typically assessed in animal models prior to human trials ^19,143–152^ – adding translational potential to our working model of the mammalian brain.

## Materials and Methods

### Human FMRI data

The dataset of functional and structural neuroimaging data used in this work came from the Human Connectome Project (HCP, http://www.humanconnectome.org/), Release Q3 ^81,153^. Per HCP protocol, all subjects gave written informed consent to the HCP consortium. These data contained fMRI and diffusion weighted imaging (DWI) acquisitions from 100 unrelated subjects of the HCP 900 data release^81,153^. All HCP scanning protocols were approved by the local Institutional Review Board at Washington University in St. Louis. Detailed information about the acquisition and imaging is provided in the dedicated HCP publications. Briefly: anatomical (T1-weighted) images were acquired in axial orientation, with FOV = 224 × 224 mm, voxel size 0.7 mm3 (isotropic), TR 2,400ms, TE 2.14ms, flip angle 8°. Functional MRI data (1200 volumes) were acquired with EPI sequence, 2 mm isotropic voxel size, TR 720ms, TE 33.1ms, flip angle 52°, 72 slices.

#### Functional MRI preprocessing and denoising

We used the minimally preprocessed fMRI data from the HCP, which includes bias field correction, functional realignment, motion correction, and spatial normalisation to Montreal Neurological Institute (MNI-152) standard space with 2mm isotropic resampling resolution. We also removed the first 10 volumes, to allow magnetisation to reach steady state. Additional denoising steps were performed using the CONN toolbox (http://www.nitrc.org/projects/conn), version 17f ^154^.

To reduce noise due to cardiac and motion artifacts, we applied the anatomical CompCor method of denoising the functional data. The anatomical CompCor method (also implemented within the CONN toolbox) involves regressing out of each individual’s functional data the first 5 principal components corresponding to white matter signal, and the first 5 components corresponding to cerebrospinal fluid signal, as well as six subject-specific realignment parameters (three translations and three rotations) and their first-order temporal derivatives, and nuisance regressors identified by the artifact detection software *art* ^155^. The subject-specific denoised BOLD signal time-series were linearly detrended and band-pass filtered between 0.008 and 0.09 Hz to eliminate both low-frequency drift effects and high-frequency noise.

Human brains were parcellated into 100 regions of interest (ROIs) covering the entire cortex. The 100 cortical ROIs were obtained from the scale-100 version of the recent local-global functional parcellation of Schaefer et al (2018) ^121^. We replicated our main results using the scale-200 Schaefer cortical atlas, augmented with augmented with 32 subcortical ROIs from a recent subcortical functional parcellation ^122^. We refer to this 232-ROI parcellation as the augmented “Schaefer-232” parcellation. The timecourses of denoised BOLD signals were averaged between all voxels belonging to a given atlas-derived ROI, using the CONN toolbox.

### Macaque FMRI data

The non-human primate MRI data were made available as part of the Primate neuroimaging Data-Exchange (PRIME-DE) monkey MRI data sharing initiative, a recently introduced open resource for non-human primate imaging ^84^.

#### Macaque dataset description

We used fMRI data from rhesus macaques (Macaca mulatta) scanned at Newcastle University. This samples includes 14 exemplars (12 male, 2 female); Age distribution: 3.9-13.14 years; Weight distribution: 7.2-18 kg (full sample description available online: http://fcon_1000.projects.nitrc.org/indi/PRIME/files/newcastle.csv and http://fcon_1000.projects.nitrc.org/indi/PRIME/newcastle.html).

Out of the 14 total animals present in the Newcastle sample, 10 had awake resting-state fMRI data; of these 10, all except the first had two scanning sessions available: to maximise our statistical power, these repeated sessions were included in the analysis. Thus, the total was 19 distinct sessions across 10 individual macaques, as in our previous publication ^101^.

##### Ethics approval

All of the animal procedures performed were approved by the UK Home Office and comply with the Animal Scientific Procedures Act (1986) on the care and use of animals in research and with the European Directive on the protection of animals used in research (2010/63/EU). We support the Animal Research Reporting of In Vivo Experiments (ARRIVE) principles on reporting animal research. All persons involved in this project were Home Office certified and the work was strictly regulated by the U.K. Home Office. Local Animal Welfare Review Body (AWERB) approval was obtained. The 3Rs principles compliance and assessment was conducted by National Centre for 3Rs (NC3Rs). Animal in Sciences Committee (UK) approval was obtained as part of the Home Office Project License approval.

##### Animal care and housing

All animals were housed and cared for in a group-housed colony, and animals performed behavioural training on various tasks for auditory and visual neuroscience. No training took place prior to MRI scanning.

#### Macaque MRI acquisition

Animals were scanned in a vertical Bruker 4.7T primate dedicated scanner, with single channel or 4-8 channel parallel imaging coils used. No contrast agent was used. Optimization of the magnetic field prior to data acquisition was performed by means of 2nd order shim, Bruker and custom scanning sequence optimisation.

Animals were scanned upright, with MRI compatible head-post or non-invasive head immobilisation, and working on tasks or at rest (here, only resting-state scans were included). Eye tracking, video and audio monitoring were employed during scanning.

Resting-state scanning was performed for 21.6 minutes, with a TR of 2600ms, 17ms TE, Effective Echo Spacing of 0.63ms, voxels size 1.22 x 1.22 x 1.24. Phase Encoding Direction: Encoded in columns. Structural scans comprised a T1 structural, MDEFT sequence with the following parameters: TE: 6ms; TR: 750 ms; Inversion delay: 700ms; Number of slices: 22; In-plane field of view: 12.8 x 9.6cm2 on a grid of 256 x 192 voxels; Voxel resolution: 0.5 x 0.5 x 2mm; Number of segments: 8.

#### Macaque functional MRI preprocessing and denoising

The macaque MRI data were preprocessed using the recently developed pipeline for non-human primate MRI analysis, *Pypreclin* (https://github.com/neurospin/pypreclin), which addresses several specificities of monkey research. The pipeline is described in detail in the associated publication ^156^. Briefly, it includes the following steps: (i) Slice-timing correction. (ii) Correction for the motion-induced, time-dependent B0 inhomogeneities. (iii) Reorientation from acquisition position to template; here, we used the recently developed National Institute of Mental Health Macaque Template (NMT): a high-resolution template of the average macaque brain generated from in vivo MRI of 31 rhesus macaques (Macaca mulatta) ^157^. (iv) Realignment to the middle volume using FSL MCFLIRT function. (v) Normalisation and masking using Joe’s Image Program (JIP) -align routine (http://www.nmr.mgh.harvard.edu/~jbm/jip/, Joe Mandeville, Massachusetts General Hospital, Harvard University, MA, USA), which is specifically designed for preclinical studies: the normalization step aligns (affine) and warps (non-linear alignment using distortion field) the anatomical data into a generic template space. (vi) B1 field correction for low-frequency intensity non-uniformities present in the data. (vii) Coregistration of functional and anatomical images, using JIP -align to register the mean functional image (moving image) to the anatomical image (fixed image) by applying a rigid transformation. The anatomical brain mask was obtained by warping the template brain mask using the deformation field previously computed during the normalization step. Then, the functional images were aligned with the template space by composing the normalization and coregistration spatial transformations.

Denoising: The aCompCor denoising method implemented in the CONN toolbox was used to denoise the macaque functional MRI data, to ensure consistency with the human data analysis pipeline. White matter and CSF masks were obtained from the corresponding probabilistic tissue maps of the high-resolution NMT template (eroded by 1 voxel); their first five principal components were regressed out of the functional data, as well as linear trends and 6 motion parameters (3 translations and 3 rotations) and their first derivatives. Finally, data were bandpass-filtered in the range of 0.008-0.09Hz Hz, as in our previous work with these data ^101^. Macaque functional data were parcellated according to the 82-ROI “Regional Mapping” cortical atlas of Kotter and Wanke ^158^, nonlinearly registered to the NMT template used for preprocessing.

### Mouse FMRI data

The mouse fMRI data used here have been previously reported ^85^. For clarity and consistency of reporting, where possible we use the same wording as in the original publication ^85^.

#### Animals and ethics

In vivo experiments were conducted in accordance with the Italian law (DL 26/214, EU 63/2010, Ministero della Sanita, Roma) and with the National Institute of Health recommendations for the care and use of laboratory animals ^85^. The animal research protocols for this study were reviewed and approved by the Italian Ministry of Health and the animal care committee of Istituto Italiano di Tecnologia (IIT). All surgeries were performed under anesthesia.

Adult (< 6 months old) male C57BL/6J mice were used throughout the study. Mice were group housed in a 12:12 hours light-dark cycle in individually ventilated cages with access to food and water ad libitum and with temperature maintained at 21 ± 1 degrees centigrade and humidity at 60 ± 10%. All the imaged mice were bred in the same vivarium and scanned with the same MRI scanner and imaging protocol employed for the awake scans (see below).

#### Experimental groups and datasets

A group of mice (n = 10, awake dataset) underwent head-post surgery, scanner habituation and fMRI image acquisitions as described below. See ^85^ for the full surgical, habituation, and scanner protocol. The scans so obtained constitute the awake rsfMRI dataset we used throughout our study.

#### MRI data acquisition

For awake scanning, the mouse was secured using an implanted headpost the custom-made MRI-compatible animal cradle and the body of the mouse was gently restrained (for details of the headpost implantation and habituation protocol, see the original publication ^85^).

All scans were acquired at the IIT laboratory in Rovereto (Italy) on a 7.0 Tesla MRI scanner (Bruker Biospin, Ettlingen) with a BGA-9 gradient set, a 72 mm birdcage transmit coil, and a four-channel solenoid receive coil. Awake rsfMRI scans were acquired using a single-shot echo planar imaging (EPI) sequence with the following parameters: TR/TE=1000/15 ms, flip angle=60 degrees, matrix=100 x 100, FOV=2.3 x 2.3 cm, 18 coronal slices (voxel-size 230 x 230 x 600 mm), slice thickness=600 mm and 1920 time points, for a total time of 32 minutes.

#### Functional MRI preprocessing, denoising, and timeseries extraction

Preprocessing of fMRI images was carried out as described in previous work ^85^. Briefly, the first 2 minutes of the time series were removed to account for thermal gradient equilibration. RsfMRI timeseries were then time despiked (3dDespike, AFNI), motion corrected (MCFLIRT, FSL), skull stripped (FAST, FSL) and spatially registered (ANTs registration suite) to an in-house mouse brain template with a spatial resolution of 0.23 x 0.23 x 0.6mm^3^. Denoising involved the regression of 25 nuisance parameters. These were: average cerebral spinal fluid signal plus 24 motion parameters determined from the 3 translation and rotation parameters estimated during motion correction, their temporal derivatives and corresponding squared regressors. No global signal regression was employed. In-scanner head motion was quantified via calculations of frame-wise displacement (FD). Average FD levels in awake conditions were comparable to those obtained in anesthetized animals (halothane) under artificial ventilation (p = 0.13, t-test) ^85^. To rule out a contribution of residual head-motion, we further introduced frame-wise fMRI scrubbing (FD > 0.075 mm). The resulting time series were band-pass filtered (0.01-0.1 Hz band) and then spatially smoothed with a Gaussian kernel of 0.5 mm full width at half maximum. The timeseries were trimmed to ensure that the same number of timepoints were included for all animals, resulting in 1414 volumes per animal. Finally data were parcellated into 72 cortical symmetric regions from the Allen Mouse Brain Atlas (CCFv3).

### Species-specific connectomes

#### Human structural connectome

We used diffusion-weighted imaging (DWI) MRI data from the Human Connectome Project. The DWI acquisition protocol is covered in detail elsewhere ^153^. The diffusion MRI scan was conducted on a Siemens 3T Skyra scanner using a 2D spin-echo single-shot multiband EPI sequence with a multi-band factor of 3 and monopolar gradient pulse. The spatial resolution was 1.25 mm isotropic. TR = 5500 ms, TE = 89.50 ms. The b-values were 1000, 2000, and 3000 s/mm2. The total number of diffusion sampling directions was 90, 90, and 90 for each of the shells in addition to 6 b0 images. We used the version of the data made available in DSI Studio-compatible format at http://brain.labsolver.org/diffusion-mri-templates/hcp-842-hcp-1021 ^159^.

We adopted previously reported procedures to reconstruct the human connectome from DWI data. The minimally-preprocessed DWI HCP data were corrected for eddy current and susceptibility artifact. After preprocessing, the DTI data were reconstructed using the model-free q-space diffeomorphic reconstruction algorithm (QSDR) implemented in DSI Studio (www.dsi-studio.labsolver.org) ^160^, following our previous work ^161^. QSDR initially reconstructs DWI data in native space, and subsequently computes values of quantitative anisotropy (QA) in each voxel, based on which DSI Studio performs a nonlinear warp from native space to a template QA volume in Montreal Neurological Institute (MNI) space. Once in MNI standard space, spin density functions are reconstructed, with a mean diffusion distance of 1.25 mm with three fiber orientations per voxel ^160^. Finally, fiber tracking was carried out by means of DSI Studio’s own “FACT” deterministic tractography algorithm, requesting 1,000,000 streamlines according to widely adopted parameters ^161^: angular cutoff = 55◦, step size = 1.0 mm, tract length between 10mm (minimum) and 400mm (maximum), no spin density function smoothing, and QA threshold determined by DWI signal in the cerebro-spinal fluid. Streamlines were automatically rejected if they presented improper termination locations, based on a white matter mask automatically generated by applying a default anisotropy threshold of 0.6 Otsu’s threshold to the anisotropy values of the spin density function ^161^.

For each individual, their structural connectome was reconstructed by drawing an edge between each pair of regions *i* and *j* if there were white matter tracts connecting the corresponding brain regions end-to-end; edge weights were quantified as the number of streamlines connecting each pair of regions.

#### Macaque structural connectome

Anatomical (structural) connectivity data were derived from the recent macaque connectome of ^83^, which combines diffusion MRI tractrography with axonal tract-tracing studies from the CoCoMac database ^162^, representing the most complete representation of the macaque connectome currently available. Structural connectivity data are expressed as a matrix in which the 82 cortical regions of interest are displayed in x-axis and y-axis. Each cell of the matrix represents the strength of the anatomical connection between any pair of cortical areas. For consistency with the human structural connectome, a symmetrised connectome was used, as in our previous work ^163^.

#### Mouse structural connectome

For the mouse structural connectome, we used a parcellated version of the high-resolution mouse connectome of Coletta *et al.* ^82^. Below, we summarise how Coletta and colleagues obtained the high-resolution mouse structural connectome.

The present mouse structural connectome is based on “high-resolution models of the mouse brain connectome (100 μm^3^) previously released by Knox and colleagues ^164^. The Knox connectome is based on 428 viral microinjection experiments in C57BL/6J male mice obtained from the Allen Mouse Brain Connectivity Atlas (http://connectivity.brain-map.org/). The connectome data were derived from imaging enhanced green fluorescent protein (eGFP)– labeled axonal projections that were then registered to the Allen Mouse Brain Atlas and aggregated according to a voxel-wise interpolation model ^164^”.

Before constructing the SC matrix, Coletta et al. ensured symmetry along the right-left axis for all the major macrostructures of the mouse brain. To this purpose, they “flipped each macrostructure (isocortex, hippocampal formation, subcortical plate, pallidum, striatum, pons, medulla, midbrain, thalamus, hypothalamus, cerebellum, and olfactory bulb) along the sagittal midline (once for the right hemisphere and once for the left hemisphere) and took the intersection with the respective nonflipped macrostructure. This procedure resulted in the removal of a set of nonsymmetric voxel (total fraction, 8.6%), the vast majority of which reside in fringe white/gray matter or cerebrospinal fluid/gray matter interfaces. The removal of these nonsymmetric voxels did not substantially affect the network structure of the resampled connectome, as assessed with a spatial correlation analysis between the symmetrized and nonsymmetrized right ipsilateral (i.e., squared) connectome”. Coletta et al. then filtered out fiber tracts and ventricular spaces, and estimated SC using a resampled version of the recently published voxel scale model of the mouse structural connectome ^164^, to make the original matrix computationally tractable. Resampling of the Knox et al. connectome was carried out by aggregating neighboring voxels according to a Voronoi diagram based on Euclidean distance between neighboring voxels, preserving the intrinsic architectural foundation of the connectome while minimizing spatial blurring and boundary effects between ontogenically distinct neuroanatomical divisions of the mouse brain, or white/gray matter, and parenchymal/ventricular interfaces (see Coletta et al. for details of the Voronoi aggregation scheme ^164^). By averaging the connectivity profile of neighboring voxels based on their relative spatial arrangement, this strategy has also the advantage of mitigating limitations related to the enforced smoothness of source space used by the original kernel interpolation used by^164^.

A whole-brain connectome was then built under the assumption of brain symmetry ^82^. Forty-four dangling nodes (i.e., nodes with no outgoing connectivity) were next removed from the resulting matrix, resulting in a final weighted and directed 15,314 × 15,314 matrix composed of 0.027-mm3 aggregate Voronoi voxels. The obtained Voronoi diagram made it possible to map the results back into the original 100-μm three-dimensional coordinate system of the Allen Institute mouse brain connectome [CCFv3]. For each pair of the 72 Allen Atlas cortical regions that we employed, their structural connectivity was obtained by averaging the connectivity of the respective constituent voxels.

To facilitate comparison of results between human and other species, the structural connectomes of macaque and mouse were symmetrised, to avoid imposing structural asymmetries and instead allow the model itself to determine the most suitable level of asymmetry (if any). Similarly, since the human and macaque structural connectomes are sparse, whereas the mouse connectome is provided as fully dense, the latter was thresholded to 50% density. However, we demonstrate that our results are not dependent on either of these methodological choices, as shown in the Supplementary Information.

### Generative whole-brain modelling

The local dynamics of each individual node is described by the normal form of a supercritical Hopf bifurcation with stochastic input, which is able to describe the transition from asynchronous noisy behaviour to full oscillations. In this regime, the Hopf model generates neither mere noise, nor the single sustained oscillation of Wilson-Cowan and Kuramoto models, but rather a fluctuating stochastically structured signal with oscillatory components that matches the infra-slow fluctuations typically observed in fMRI signal ^165^.

Thus, each node *n* is represented by the following set of coupled stochastic differential equations in Cartesian coordinates:

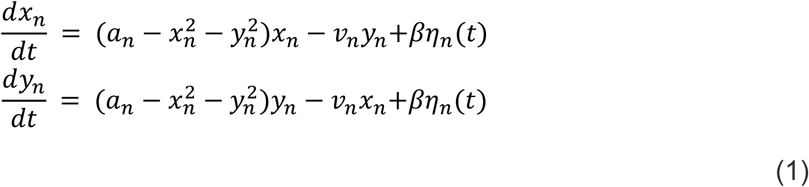

In these equations, *η* represents additive Gaussian noise with a standard deviation*β*. The system undergoes a supercritical bifurcation at *a*_*n*_ = 0. When *a*_*n*_ > 0, the system engages in a stable limit cycle with a frequency 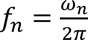, while for *a*_n_ < 0, the dynamics stabilise at a fixed point, representing a low-activity noisy state dominated by the Gaussian noise. In this model, each node has an intrinsic frequency *ω*_*n*_ within the range of the empirical fMRI range, determined by the averaged peak frequency of the narrowband BOLD signals in each brain region. We set *a*_*n*_ = −0.02, consistent with previous studies ^78^.

To model whole-brain dynamics, we incorporate coupling between regions using a diffusive coupling term. This term represents the input received by region *n* from every other region *p,* weighted by the corresponding connection from the adjacency matrix, *G*_*np*_ representing empirical structural connectivity. The input is modelled using a common difference coupling, approximating the simplest (linear) component of a general coupling function. The equations governing the whole-brain dynamics are:

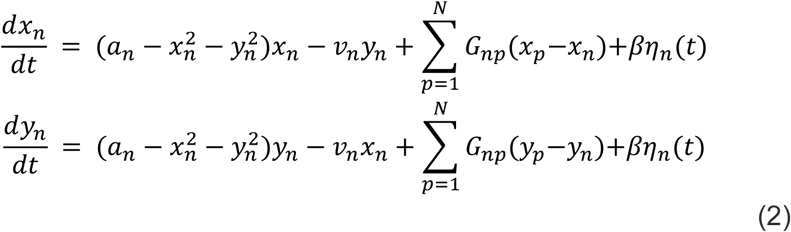

In these equations, the noise standard deviation is set to *β* = 0.01. This coupled oscillator model effectively captures the transition between different dynamical states of the brain and provides a framework for understanding how brain regions interact to produce complex patterns of activity.

#### Updating the Generative Effective Connectivity

We optimised the generative effective connectivity (GEC) between brain regions by aligning the model’s output with empirical measures, specifically forward and reverse time-shifted correlations, and empirical functional connectivity (FC). A heuristic gradient algorithm was then employed to iteratively update the GEC, refining the fit ^78^.

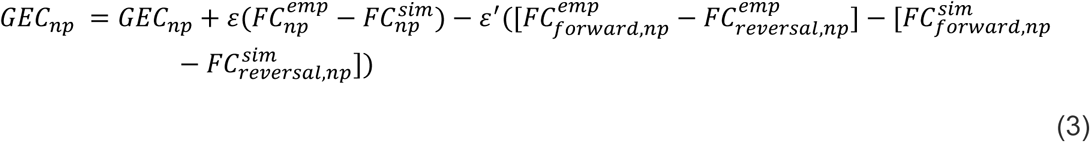

The model was iteratively run with the updated GEC until the fit converged to a stable value.

Initialization was based on anatomical connectivity, with updates applied only to known existing connections within the matrix. Following previous work (REF), the algorithm was run with ε = 0.0002 and ε′ = 0.00004, continuing until convergence was achieved. Crucially, two versions of the algorithm were considered: *cooperative-only,* whereby connection updates that reached negative values were not allowed (as is current practice); and *cooperative+competitive,* whereby the algorithm is allowed to update the connections to negative values.

### Differential identifiability for model evaluation

“Brain fingerprinting” refers to using brain-derived metrics (here: the functional connectivity obtained from resting-state functional MRI) to discriminate individuals from each other, analogously to how the grooves on one’s fingertips may be used to discern one’s identity ^87,166^. This requires brain fingerprints (just like conventional fingerprints) to be different across different people (to avoid confusing distinct individuals) but consistent within the same individual (to track identity).

Let A be the “identifiability matrix”, i.e. the of similarity between individuals’ test and retest scans (or in this case, individuals’ empirical and simulated FC), such that the size of A is S-by-S (with S being the number of individuals in the dataset). Each entry of A is obtained as the correlation between the corresponding individuals’ vectorised matrices of functional connectivity. Let I_self_ = ⟨A_ii_⟩ represent the average of the main diagonal elements of A, which consist of the Pearson correlation values between scans of same individual: from now on, we will refer to this quantity as self-identifiability or I_self_. Similarly, let I_others_ = ⟨A_ij_⟩ define the average of the off-diagonal elements of matrix A, i.e. the correlation between models and scans of different individuals i and j. Then, we define the differential identifiability (I_diff_) of the sample as the difference between both terms ^87^:

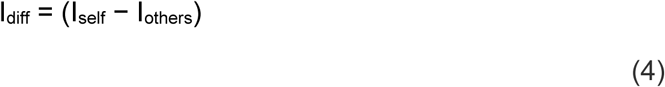

which quantifies the difference between the average FCs similarity using the matched model, and the average FCs similarity when model and empirical data are mismatched. The higher the value of I_diff_, the higher the model’s individual-specificity ^87,110^.

### Dynamical measures

#### Synchrony and Metastability

Metastability was quantified using a widely-used signature, the standard deviation of the Kuramoto Order Parameter across time (*std*(*KOP*)) ^77,94,95^.

In turn, the Kuramoto Order Parameter is defined by following equation:

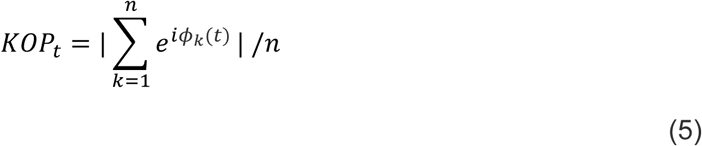

where *φ_k_*(*t*) is the instantaneous phase of each bandpass-filtered BOLD signal at node *k*. Following ^77^: “we computed the instantaneous phase φ_k_(t) of each bandpass-filtered signal *k* using the Hilbert transform. The Hilbert transform yields the associated analytical signals. The analytic signal represents a narrowband signal, s(t), in the time domain as a rotating vector with an instantaneous phase, φ(t), and an instantaneous amplitude, A(t). Thus, *s*(*t*) = *A*(*t*)cos(*ϕ*(*t*)). The phase and the amplitude are given by the argument and the modulus, respectively, of the complex signal *z*(*t*), given by *z*(*t*)=*s*(*t*)+*iH*[*s*(*t*)], where *i* is the imaginary unit and H[*s*(*t*)] is the Hilbert transform of *s*(t)”. At each point in time, the Kuramoto order parameter measures the global level of synchronization across these oscillating signals. Under complete independence, the phases are uniformly distributed and thus KOP is nearly zero, whereas KOP = 1 if all phases are equal (full synchronization) ^167^.

The variability (standard deviation) of the KOP over time is commonly used as a signature of metastability, first proposed by Shanahan ^77,94,95,168^. See ^95^ for a detailed discussion of the difference between metastability and its various signatures. Intuitively, if *std(KOP)* is high, it indicates that the system alternates between high and low synchronisation, thereby combining tendencies for integration (high synchrony) and segregation (low synchrony). For each individual (and simulation), we obtain the *std(KOP)* signature of metastability, as well as the maximum observed value of global synchrony.

#### Local-global hierarchy from Intrinsic-driven Ignition

“Intrinsic-driven ignition” ^97^ quantifies the extent to which spontaneously occurring (“intrinsic”) neural events elicit activation across the rest of the brain (“ignition”). The fMRI signal timeseries are transformed into z-scores, and subsequently thresholded to obtain a binary sequence σ based on the combined mean and standard deviation of the regional transformed signal, such that σ(t) = 1 if z(t) > 1 and is crossing the threshold from below, indicating that a local event has been triggered; otherwise, σ(t) = 0. Subsequently, for each brain region, when that region triggers a local event (σ(t) = 1, the resulting global ignition is computed within a time-window of 4 TRs (note that the threshold of 1 standard deviation and window of 4 TRs for defining an event are chosen for consistency with previous work ^45,97^, but it has been demonstrated that the results of this procedure are robust to the specific threshold chosen ^169^).

An NxN binary matrix M is then constructed, indicating whether in the period of time under consideration two regions *i* and *j* both triggered an event (M*_ij_ =* 1). The size of the largest connected component of this binary matrix M defines the breadth of the global ignition generated by the driver region at time *t,* termed “intrinsic-driven ignition” (IDI). To obtain a measure of local-global hierarchy, the variability (standard deviation) across event sizes is then computed ^45,97^. Consequently, higher standard deviation reflects more heterogeneity with respect to regions’ capability to induce ignition, which suggests in turn a more elaborate hierarchical organisation between them.

#### Hierarchy from temporal irreversibility

We estimate pairwise interactions between brain regions by computing time-shifted correlations between both the forward and reversed fMRI BOLD time series of any two regions ^78,98^. This method effectively quantifies the asymmetry in interactions between region pairs, thereby indicating how one region influences another. This approach is inspired by thermodynamics, where the breaking of detailed balance is associated with non-reversibility, often referred to as the “arrow of time”. Irreversibility is captured as the difference between the time-shifted correlations of forward and reverse time series ^78,98^.

To illustrate, consider the detection of irreversibility between two time series, x(t) and y(t). The causal dependency between x(t) and y(t) is measured using time-shifted correlations. For forward evolution, the time-shifted correlation is given by:

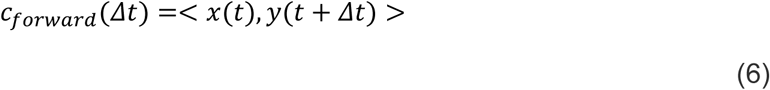

Similarly, we create a reversed version of x(t) (or y(t)), denoted *x*^*r*^(*t*) (or *y*^*r*^(*t*)), by inverting the time sequence. The time-shifted correlation for the reversed evolution is then:

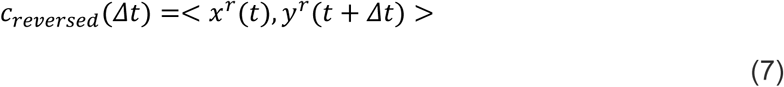

The pairwise level of irreversibility, representing the degree of temporal asymmetry or the arrow of time, is quantified as the absolute difference between the forward and reversed time-shifted correlations at a given shift *Δ*t=T (here, for consistency across species we set T=1 TR):

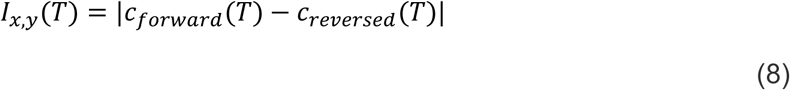

To compute the whole-brain level of NR, we defined forward and reversal matrices of time-shifted correlations for the forward version *x*_*n*_ and the reversed backward version 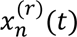 of a multidimensional time series, where the subscript *n* represents different brain regions. These matrices capture the functional causal dependencies between the variables in the forward and artificially generated reversed time series, respectively. The forward and reversed matrices are expressed as:

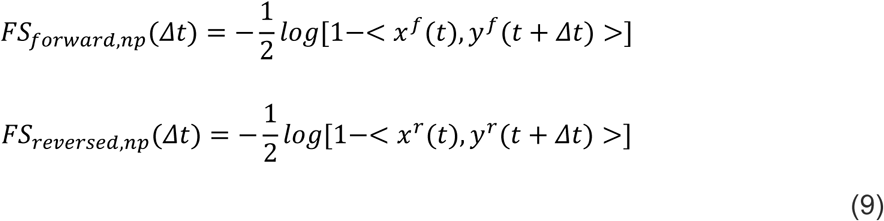

These matrices, representing the functional temporal dependencies, are based on the mutual information derived from the respective time-shifted correlations ^78,98^.

*FS*_*diff*,*np*_ is a matrix containing the squared differences of the elements between the forward and reversed matrices:

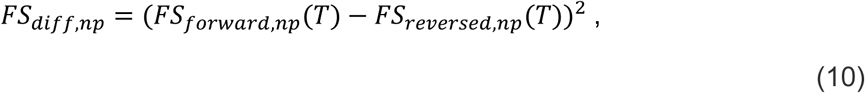

where each element reflects the irreversibility level for that region pair. We describe the level of hierarchy as the standard deviation of the elements of the matrix *FS*_*diff*,*np*_ ^78,98^.

Irreversibility between two regions occurs when the interaction between them is asymmetric, such that one region sends more information to the other than it receives from it ^78,98^. For each region, its mean irreversibility with the rest of the brain therefore quantifies how far it is from equilibrium between sending and receiving. The variability across regions of this send-receive imbalance provides a measure of hierarchical organisation, analogous to the ignition-based hierarchy. If this variability is high, it means that regions vary widely in their preference for sending or receiving signals, reflecting greater hierarchical character (in terms of directedness, rather than local-global ignition) of the functional organisation ^78,98^.

#### Synergistic information

The framework of integrated information decomposition (ΦID) unifies integrated information theory (IIT) and partial information decomposition (PID) to decompose information flow into interpretable, disjoint parts. In this section we provide a brief description of ΦID and formulae required to compute the results. For further details, see ^101,102^.

##### Partial information decomposition

We begin with Shannon’s Mutual information (MI), which quantifies the interdependence between two random variables *X* and *Y*. It is calculated as

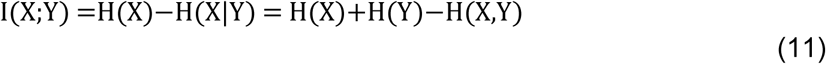

where H(*X*) stands for the Shannon entropy of a variable *X*. Above, the first equality states that the mutual information is equal to the reduction in entropy (i.e., uncertainty) about *X* after *Y* is known. Put simply, the mutual information quantifies the information that one variable provides about another ^170^.

Crucially, Williams and Beer ^106^ observed that the information that two source variables *X* and *Y* give about a third target variable *Z*, I(*X,Y*; *Z*), should be decomposable in terms of different *types* of information: information provided by one source but not the other (unique information), by both sources separately (redundant information), or jointly by their combination (synergistic information; Figure 2A). Following this intuition, they developed the *Partial Information Decomposition* (PID; ^106^) framework, which leads to the following fundamental decomposition:

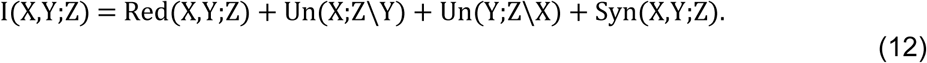

Above, *Un* corresponds to the unique information one source but the other doesn’t, *Red* is the redundancy between both sources, and *Syn* is their synergy: information that neither *X* nor *Y* alone can provide, but that can be obtained by considering *X* and *Y* together.

The simplest example of a purely synergistic system is one in which *X* and *Y* are independent fair coins, and *Z* is determined by the exclusive-OR function *Z* = XOR(*X,Y*): i.e., *Z*=0 whenever *X* and *Y* have the same value, and *Z*=1 otherwise. It can be shown that *X* and *Y* are both statistically independent of *Z*, which implies that neither of them provide - by themselves - information about *Z*. However, *X* and *Y* together fully determine *Z*, hence the relationship between *Z* with *X* and *Y* is purely synergistic.

As another example for the case of Gaussian variables (as employed here), consider a 2-node coupled autoregressive process with two parameters: a noise correlation *c* and a coupling parameter *a*. As *c* increases, the system is flooded by “common noise”, making the system increasingly redundant because the common noise “swamps” the signal of each node. As *a* increases, each node has a stronger influence both on the other and on the system as a whole, and we expect synergy to increase. Therefore, synergy reflects the joint contribution of parts of the system to the whole that is not driven by common noise. This can be demonstrated empirically ^102,107^.

##### Synergy from Integrated Information Decomposition

Recently, Mediano, Rosas and colleagues ^102^ formulated an extension of PID able to decompose the information that multiple source variables have about multiple target variables. This makes PID applicable to the dynamical systems setting, and yields a decomposition with redundant, unique, and synergistic components in the past and future that can be used as a principled method to analyse information flow in neural activity.

While there is ongoing research on the advantages of different information decompositions for discrete data, several decompositions converge into the same simple form for the case of univariate Gaussian variables ^171^. Known as *minimum mutual information PID* (MMI-PID), this decomposition quantifies redundancy in terms of the minimum mutual information of each individual source with the target; synergy, then, becomes identified with the additional information provided by the weaker source once the stronger source is known. Since linear-Gaussian models are sufficiently good descriptors of functional MRI timeseries (and more complex, non-linear models have been shown to offer no significant advantage ^172,173^), here we adopt the MMI-PID decomposition, following our own and others’ previous applications of PID to neuroscientific data ^101,102^.

In a dynamical system such as the brain, one can calculate the amount of information flowing from the system’s past to its future, known as time-delayed mutual information (TDMI). Specifically, by denoting the past of variables as *X_t-τ_* and *Y_t-τ_* and treating them as sources, and their joint future state (*X_t_*, *Y_t_*), as target, one can apply the PID framework and decompose the information flowing from past to future as

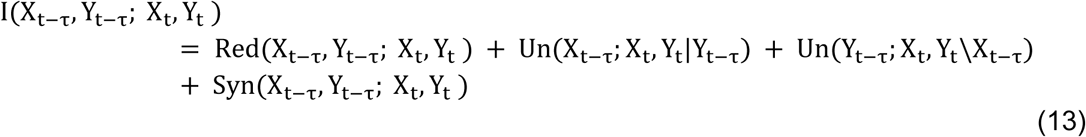

Applying ΦID to this quantity allows us to distinguish between redundant, unique, and synergistic information shared with respect to the future variables *X_t_*, *Y_t_* ^101,102^. Importantly, this framework has identified a stronger notion of redundancy, in which information is shared by *X* and *Y* in both past and future. Accordingly, using the MMI-ΦID decomposition for Gaussian variables, we use

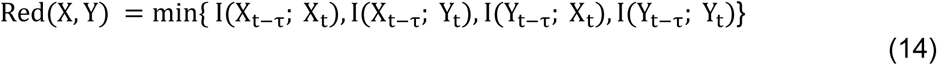

Equivalently, this measure corresponds to the information that was redundantly carried by *X* and *Y* in the past, and is also redundantly carried redundantly by the two sources in the future. Using this definition of redundancy we can then solve the system of equations and recover how each type of information (synergistic, unique, and redundant) evolves over time. Of these, we focus on the temporally persistent synergy (denoted by 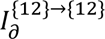 in standard PID notation): information that was synergistic in the past, and remains synergistic in the future). We have released MATLAB/Octave and Python code to compute measures of Integrated Information Decomposition of timeseries with the Gaussian MMI solver, available at https://github.com/Imperial-MIND-lab/integrated-info-decomp.

### Cognitive matching from NeuroSynth

We recently introduced “cognitive matching” in ^110^. For consistency, we report this procedure using the same wording as in ^110^: “Continuous measures of the association between voxels and cognitive categories were obtained from NeuroSynth, an automated term-based meta-analytic tool that synthesizes results from more than 14,000 published fMRI studies by searching for high-frequency key words (such as “pain” and “attention” terms) that are systematically mentioned in the papers alongside fMRI voxel coordinates (https://github.com/neurosynth/neurosynth), using the volumetric association test maps ^115^. This measure of association strength is the tendency that a given term is reported in the functional neuroimaging study if there is activation observed at a given voxel. Note that NeuroSynth does not distinguish between areas that are activated or deactivated in relation to the term of interest, nor the degree of activation, only that certain brain areas are frequently reported in conjunction with certain words.

Although more than a thousand terms are catalogued in the NeuroSynth engine, we refine our analysis by focusing on cognitive function and therefore we limit the terms of interest to cognitive and behavioural terms. To avoid introducing a selection bias, we opted for selecting terms in a data-driven fashion instead of selecting terms manually. Therefore, terms were selected from the Cognitive Atlas, a public ontology of cognitive science ^114^, which includes a comprehensive list of neurocognitive terms. This approach totaled to t = 123 terms, ranging from umbrella terms (“attention”, “emotion”) to specific cognitive processes (“visual attention”, “episodic memory”), behaviours (“eating”, “sleep”), and emotional states (“fear”, “anxiety”) (note that the 123 term-based meta-analytic maps from NeuroSynth do not explicitly exclude patient studies). The Cognitive Atlas subdivision has previously been used in conjunction with NeuroSynth ^111–113,174^, so we opted for the same approach to make our results comparable to previous reports. The full list of terms included in the present analysis is shown in Figure S13. The probabilistic measure reported by NeuroSynth can be interpreted as a quantitative representation of how regional fluctuations in activity are related to psychological processes. As with the resting-state BOLD data, voxelwise NeuroSynth maps were parcellated into 100 cortical regions according to the Schaefer atlas (or 232 cortical and subcortical regions for the replication with subcortex included).”

For each individual, their parcellated BOLD signals at each point in time were spatially correlated against each NeuroSynth map, producing one value of correlation per NeuroSynth map, per BOLD volume. We refer to this operation as “cognitive matching”. For each volume, the quality of cognitive matching was quantified as the highest value of (positive) correlation across all maps. These values were subsequently averaged across all volumes to obtain a single value per condition, per participant / simulation.

### Computational capacity from reservoir computing

The reservoir computing architecture used in this study consists of a nonlinear recurrent neural network (RNN; reservoir) complemented by a linear readout module that approximates a target signal by means of a linear combination of the signals of output nodes selected from the reservoir ^175^. Only the readout module is trained, allowing us to constrain the reservoir with generative effective connectivity patterns that remain unchanged throughout learning. In this study, the reservoir states obey the following discrete-time update equation:

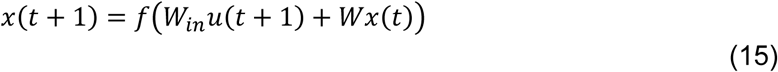

Where *x*(*t*) is the vector of nodal reservoir activation states at time *t*, *u*(*t*) is the K-dimensional input signal at time *t*, *W*_*in*_ is the input matrix mapping the input signal to the input nodes, *W* is the reservoir weight matrix, and *f* is the hyperbolic tangent. An input gain of 0.0001 was used for *W*_*in*_ following ^119^.

The readout module is trained following Ridge regression as implemented in *sklearn* using the default *α* = 0.5 l2 regularization parameter from the freely available *conn2res* package (https://github.com/netneurolab/conn2res) ^118^.

To evaluate computational capacity, we chose the widely used memory capacity task, which measures the reservoir’s ability to encode past stimuli ^117,119,176–179^. In this task, the readout module is trained to reproduce a time-delayed version of a random uniformly distributed input signal *u*(*t*)∼*U*(−1, 1). Specifically, *y*(*t*) = *u*(*t* − *τ*), where *y*(*t*) is the target signal at time *t* and *τ* is the time lag considered. For each time lag, performance at the task was evaluated as the absolute value of the Pearson correlation coefficient between the target signal *y*(*t*) and the predicted signal *y*^(*t*) obtained from the trained readout module. Memory capacity was then evaluated as the sum of the performance scores across all time lags:

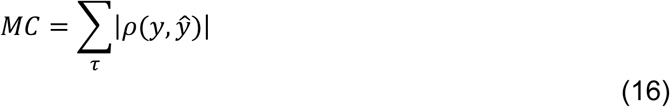

We generated 4050 input signal timepoints and used a 70:30 train/test split ratio. Reservoir states were then simulated separately for the training and testing input timeseries. The first 50 timepoints of both resulting reservoir states timeseries were discarded to account for initial transients. Memory capacity was evaluated only in the test timeseries across 20 time lags, monotonically increased in one timepoint steps in the range [1,20].

For each species, input nodes were chosen to coincide with visual cortical regions, and the somatomotor regions were used as output nodes, to reflect their respective functional roles ^118,178^. For each individual generative effective connectivity network, we averaged memory capacity across 10 simulations, selecting half of the visual regions (*K*) as input nodes at random each time. Furthermore, the connectivity matrices were normalized by their spectral radius and the connection weights were uniformly scaled to produce a range of spectral radii *α* ∈ [0.1, 1.6] in 0.1 increments. Specifically, the reservoir matrix *W* can be expressed as:

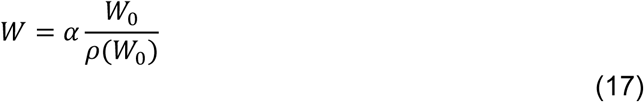

Where *W*_0_ is the original effective connectivity network and *ρ*(*W*_0_) is its spectral radius. This allows us to parametrically tune the reservoir’s global dynamics ^119,177^. For each network, only the best performance across all *α* values was retained.

### Network properties

#### Modularity

The network modularity function quantifies the extent to which a network can be partitioned such that the number of within-group edges is maximised and the density of between-group edges is minimised. We employed an implementation of Newman’s spectral modularity algorithm available in the *Brain Connectivity Toolbox* (BCT ^180,181^).

#### Clustering coefficient

The clustering coefficient of node *i* (*C_i_*) is a node-specific measure of how well connected a node’s neighbourhood is; it is calculated as the fraction of neighbours of the node that are also neighbours of each other

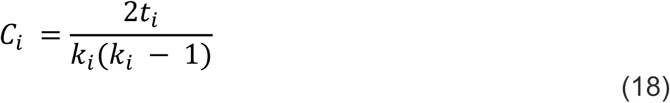

where *t_i_* is the number of triangles around node *i,* and *k*_*i*_ is the number of edges connected to node *i.* An overall measure of clustering for the entire network is obtained by averaging the nodes’ clustering coefficients. We used the implementation of clustering coefficient available _in BCT 180,181._

### Statistical reporting

Statistical significance was assessed using a resampling-based, paired-samples t-test. This non-parametric implementation of the test ensures robustness to violations of the normality assumption, which was not formally tested. Effect sizes are provided as Hedge’s measure of standardised difference *g,* which is analogous to Cohen’s *d,* but recommended for smaller sample sizes such as the ones available in the present study.

## Data availability

The Human Connectome Project functional and structural datasets are freely available from http://www.humanconnectome.org/. Macaque functional MRI data are available from the PRIMatE Data Exchange (PRIME-DE) through the Neuroimaging Informatics Tools and Resources Clearinghouse (NITRC; http://fcon_1000.projects.nitrc.org/indi/indiPRIME.html). The macaque connectome is available on Zenodo at https://doi.org/10.5281/zenodo.1471588. The CoCoMac database on which it is based, is also available online at http://cocomac.g-node.org/main/index.php?. Mouse functional and structural connectome data are available from author A.G. NeuroSynth is available at https://neurosynth.org/.

## Acknowledgements

A.I.L. was supported by St John’s College, Cambridge and the Wellcome Trust (grant number 226924/Z/23/Z). J.V. is supported by EU H2020 FET Proactive project Neurotwin (101017716). Y.S.P. is supported by European Union’s Horizon 2020 research and innovation programme under the Marie Sklodowska-Curie grant 896354, and ‘ERDF A way of making Europe’, ERDF, EU, Project NEurological MEchanismS of Injury, and Sleep-like cellular dynamics (NEMESIS; ref. 101071900) funded by the EU ERC Synergy Horizon Europe. M.L.K. is supported by the Center for Music in the Brain, funded by the Danish National Research Foundation (DNRF117), and Centre for Eudaimonia and Human Flourishing at Linacre College funded by the Pettit and Carlsberg Foundations. G.D. is supported by grant no. PID2022-136216NB-I00 funded by MICIU/AEI/10.13039/501100011033 and by ‘ERDF A way of making Europe’, ERDF, EU, Project NEurological MEchanismS of Injury, and Sleep-like cellular dynamics (NEMESIS; ref. 101071900) funded by the EU ERC Synergy Horizon Europe, and AGAUR research support grant (ref. 2021 SGR 00917) funded by the Department of Research and Universities of the Generalitat of Catalunya. A.G. is supported by Simons Foundation Grants (SFARI 400101); the European Research Council (ERC—DISCONN, No. 802371); Brain and Behavior Foundation 2017 (NARSAD—National Alliance for Research on Schizophrenia and Depression), NIH (1R21MH116473-01A1), and the Telethon Foundation (GGP19177). B.M. acknowledges support from the NSERC, Canadian Institutes of Health Research (CIHR), Brain Canada Foundation Future Leaders Fund, the Canada Research Chairs Program, the Michael J. Fox Foundation, and the Healthy Brains for Healthy Lives initiative. For the purpose of open access, the authors have applied a creative commons attribution (CC BY) licence to any author accepted version arising from this manuscript.

## Competing interests

The authors declare no competing interests.

## Notes

### Competing Interest Statement

The authors have declared no competing interest.

