## Supplementary Figures for "Competitive interactions shape brain dynamics and computation across species"

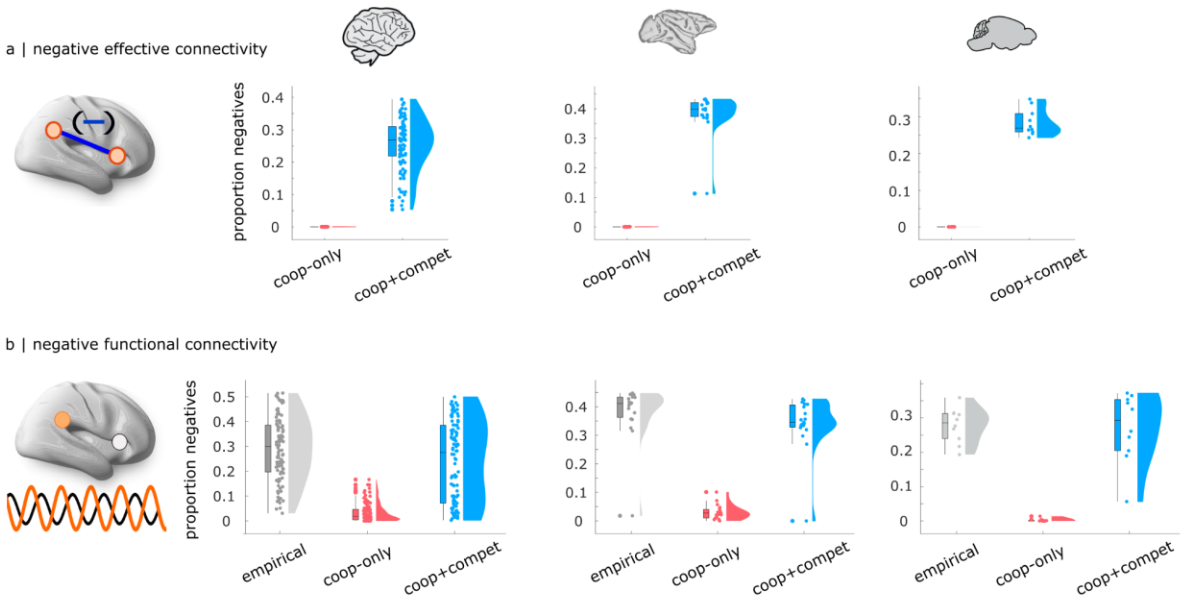


**Figure S1. Prevalence of competitive interactions in the effective connectivity and negative functional connectivity.** (**a**) As expected, the model that only allows cooperative interactions in the effective connectivity does not have any competitive interactions. In contrast, when the model is allowed to have both cooperative and competitive effective connectivity, we observe a non-zero proportion of competitive effective connections, in each individual of each species. (**b**) Individual brains vary in the prevalence of anti-correlations in the empirical FC (though always non-zero). Even the model that only allows cooperative effective connectivity can produce some negative functional connectivity. However, the proportion of negative edges in the simulated FC is significantly higher, and closer to the empirical, when competitive effective connectivity is also allowed in the model. See Supplementary Tables 1-3 for full statistical reporting.

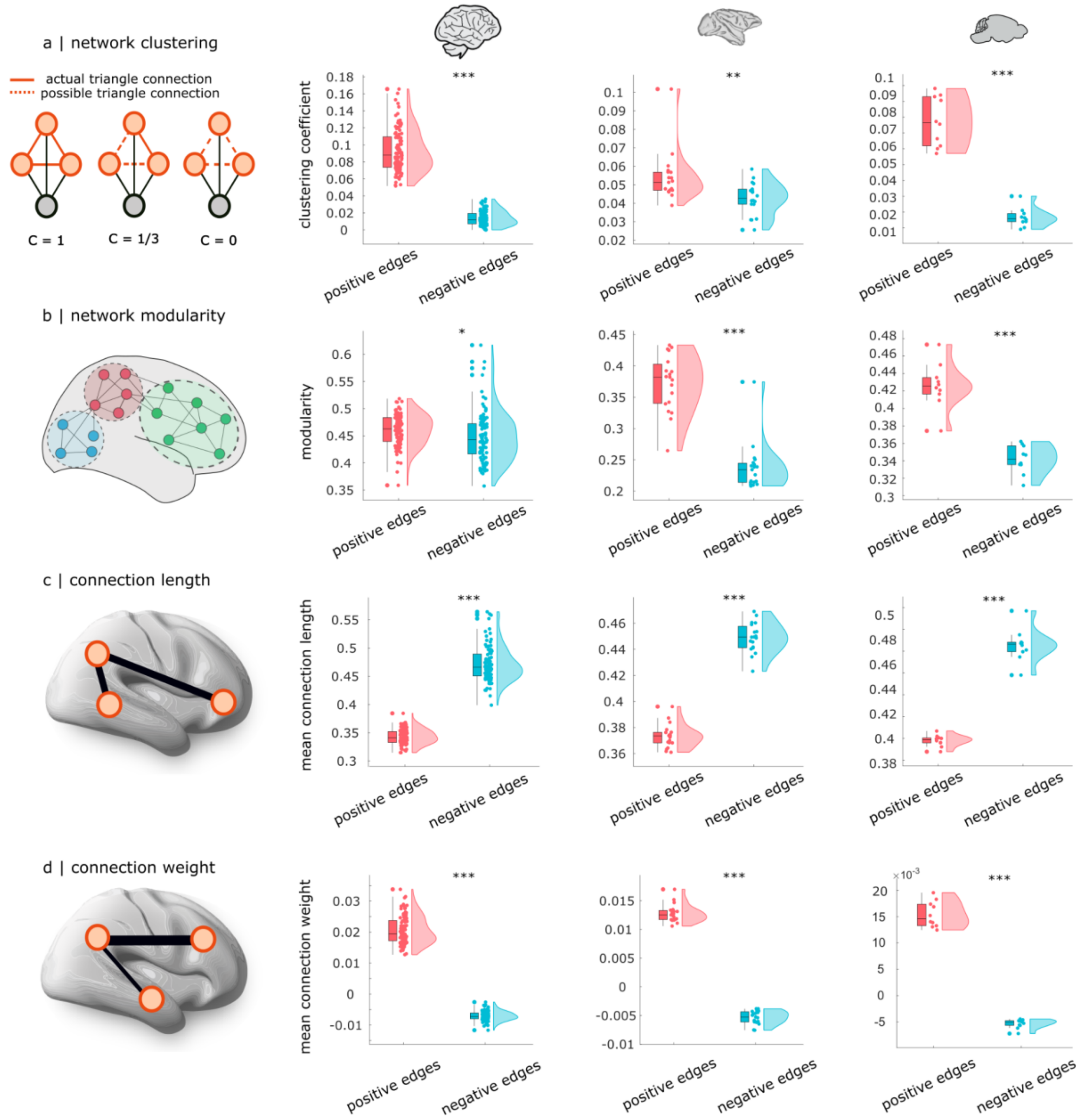


**Figure S2. Network properties of the cooperative and competitive interactions in the generative effective connectivity.** Competitive connections are significantly less clustered (**a**) and modular (**b**) than cooperative econnections. Competitive connections are also longer, connecting regions that are further apart from each other in space (**c**) but tend to be weaker in weight (**d**). For (**c**), length is the Euclidean distance between region centroids; for ease of comparison across species, the Y axis is normalised by the maximum distance. See Supplementary Tables 1-3 for full statistical reporting.

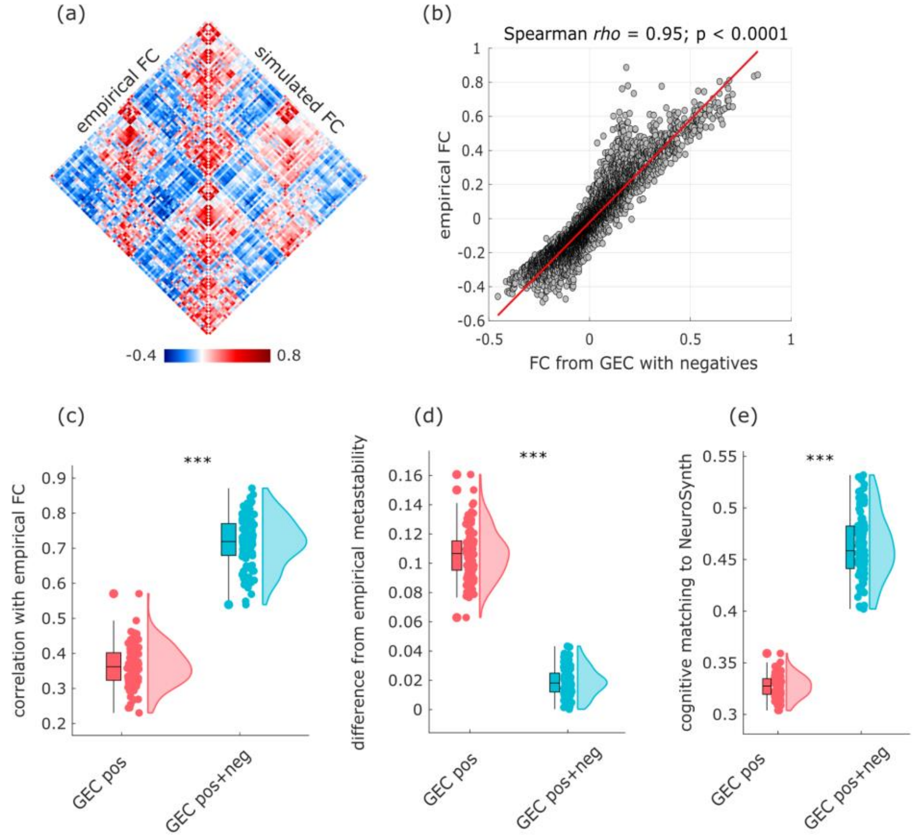


**Figure S3. Replication of human results with global signal regression.** (**a**) Group-wise empirical and simulated FC. (**b**) Correlation between group-wise empirical and simulated FC ($r$ = 0.95). (**c**) Subject-wise correlations between empirical and simulated FC are significantly higher for the model including both cooperative and competitive interactions in the effective connectivity. (**d**) Subject-wise difference between empirical and simulated metastability is significantly lower (reflecting greater similarity) for the model including both cooperative and competitive interactions in the effective connectivity. (**e**) Subject-wise cognitive matching to NeuroSynth meta-analytic maps is significantly higher for the model including both cooperative and competitive interactions in the effective connectivity.

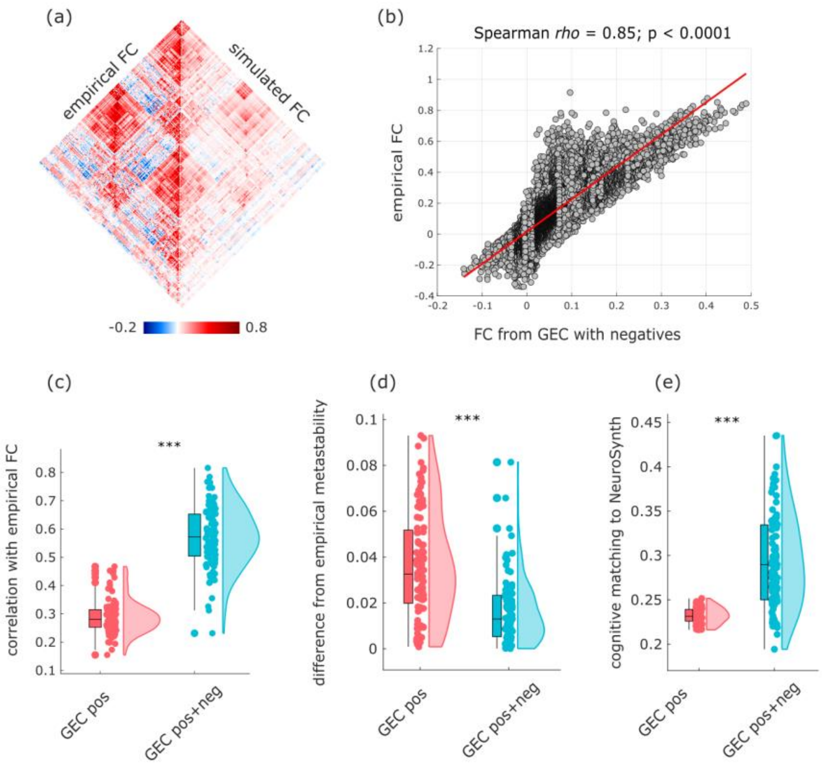


**Figure S4. Replication of human results with a different cortical parcellation including subcortex (Schaefer-232).** (**a**) Group-wise empirical and simulated FC. (**b**) Correlation between group-wise empirical and simulated FC ($r$ = 0.85). (**c**) Subject-wise correlations between empirical and simulated FC are significantly higher for the model including both cooperative and competitive interactions in the effective connectivity. (**d**) Subject-wise difference between empirical and simulated metastability is significantly lower (reflecting greater similarity) for the model including both cooperative and competitive interactions in the effective connectivity. (**e**) Subject-wise cognitive matching to NeuroSynth meta-analytic maps is significantly higher for the model including both cooperative and competitive interactions in the effective connectivity.

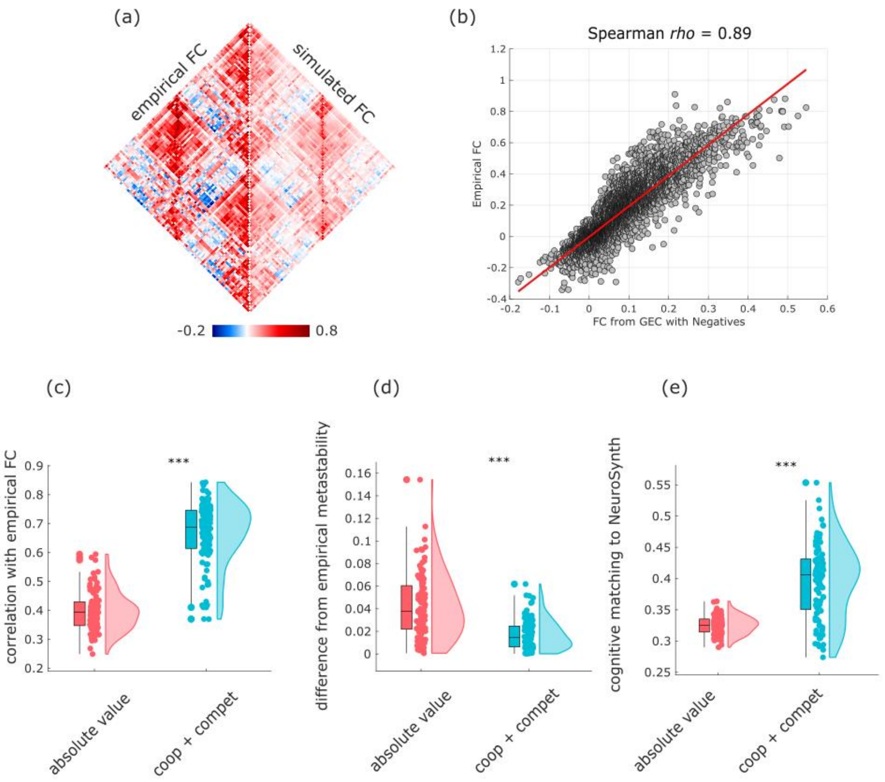


**Figure S5. Replication of human results with homotopic connections added to the SC.** (**a**) Group-wise empirical and simulated FC. (**b**) Correlation between group-wise empirical and simulated FC ($r$ = 0.89). (**c**) Subject-wise correlations between empirical and simulated FC are significantly higher for the model including both cooperative and competitive interactions in the effective connectivity. (**d**) Subject-wise difference between empirical and simulated metastability is significantly lower (reflecting greater similarity) for the model including both cooperative and competitive interactions in the effective connectivity. (**e**) Subject-wise cognitive matching to NeuroSynth meta-analytic maps is significantly higher for the model including both cooperative and competitive interactions in the effective connectivity.

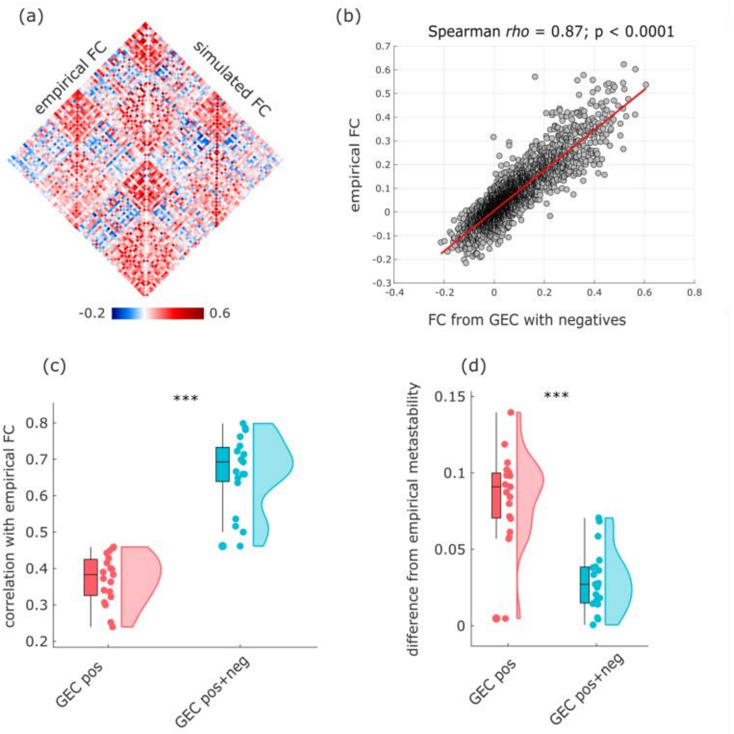


**Figure S6. Replication of macaque results with asymmetric connectome.** (**a**) Group-wise empirical and simulated FC. (**b**) Correlation between group-wise empirical and simulated FC ($r$ = 0.87). (**c**) Subject-wise correlations between empirical and simulated FC are significantly higher for the model including both cooperative and competitive interactions in the effective connectivity. (**d**) Subject-wise difference between empirical and simulated metastability is significantly lower (reflecting greater similarity) for the model including both cooperative and competitive interactions in the effective connectivity.

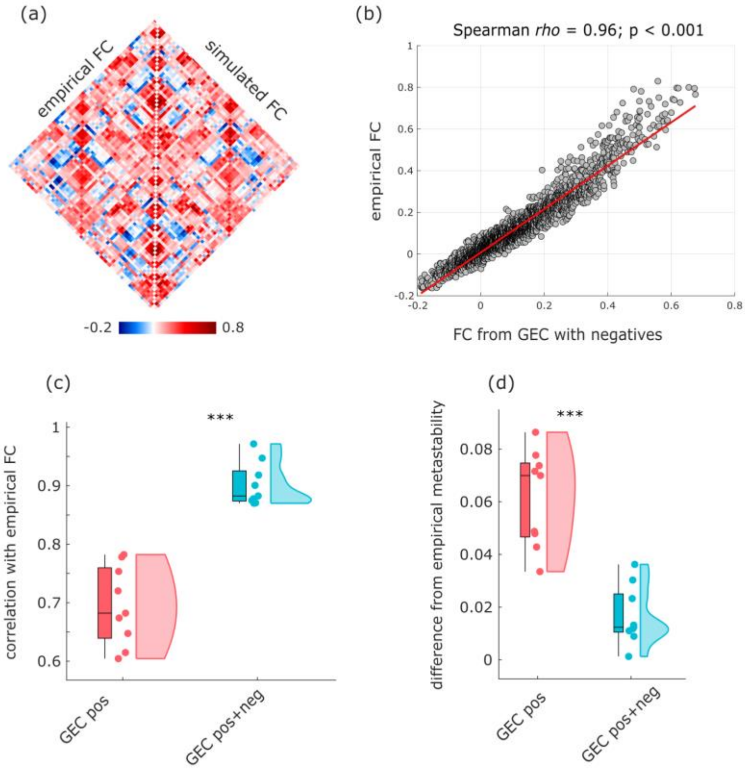

**Figure S7. Replication of mouse results with asymmetric, fully dense connectome.** (**a**) Group-wise empirical and simulated FC. (**b**) Correlation between group-wise empirical and simulated FC (*r* = 0.96). (**c**) Subject-wise correlations between empirical and simulated FC are significantly higher for the model including both cooperative and competitive interactions in the effective connectivity. (**d**) Subject-wise difference between empirical and simulated metastability is significantly lower (reflecting greater similarity) for the model including both cooperative and competitive interactions in the effective connectivity.

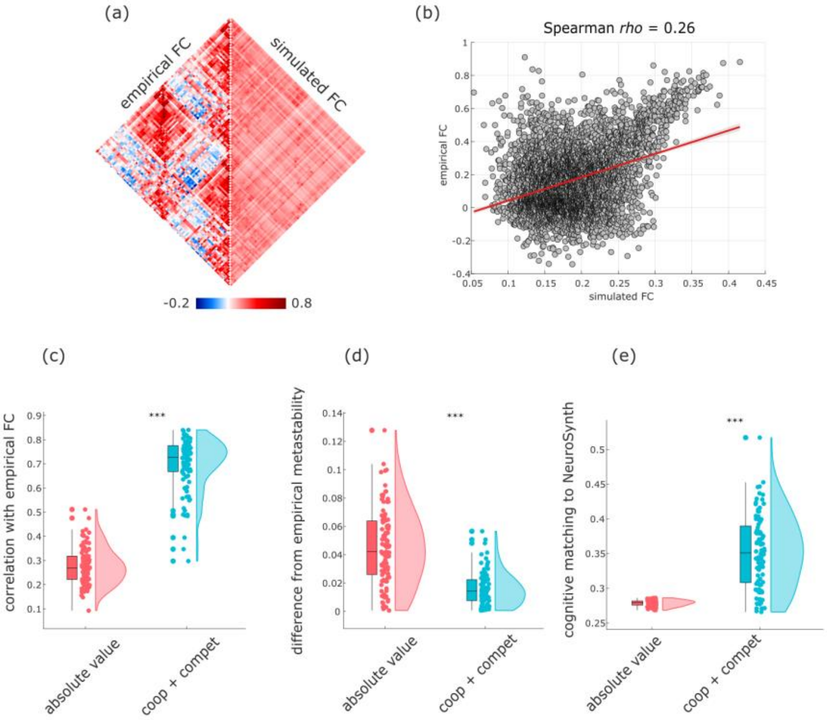


**Figure S8. Turning competitive interactions into cooperative degrades performance of human model.** (**a**) Group-wise empirical and simulated FC. (**b**) Correlation between group-wise empirical and simulated FC (*r* = 0.26). (**c**) Subject-wise correlations between empirical and simulated FC are significantly higher for the model including both cooperative and competitive interactions in the effective connectivity, than for the model where competitive interactions are turned into cooperative. (**d**) Subject-wise difference between empirical and simulated metastability is significantly lower (reflecting greater similarity) for the model including both cooperative and competitive interactions in the effective connectivity, than for the model where competitive interactions are turned into cooperative. (**e**) Subject-wise cognitive matching to NeuroSynth meta-analytic maps is significantly higher for the model including both cooperative and competitive interactions in the effective connectivity, than for the model where competitive interactions are turned into cooperative.

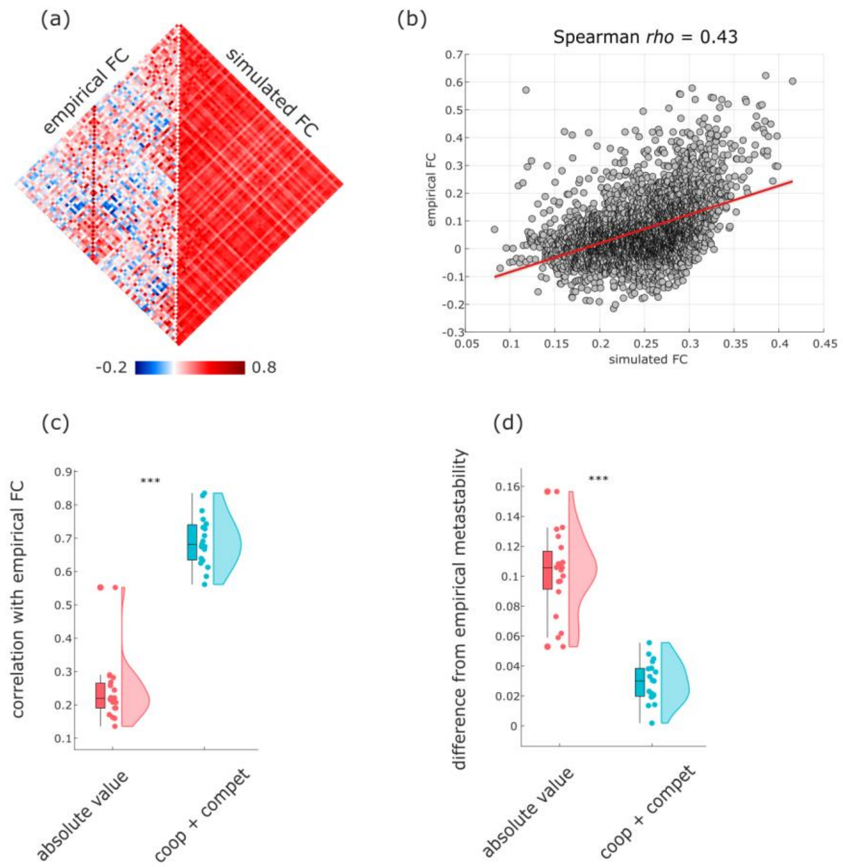


**Figure S9. Turning competitive interactions into cooperative degrades performance of macaque model.** (**a**) Group-wise empirical and simulated FC. (**b**) Correlation between group-wise empirical and simulated FC ($r = 0.43$). (**c**) Subject-wise correlations between empirical and simulated FC are significantly higher for the model including both cooperative and competitive interactions in the effective connectivity, than for the model where competitive interactions are turned into cooperative. (**d**) Subject-wise difference between empirical and simulated metastability is significantly lower (reflecting greater similarity) for the model including both cooperative and competitive interactions in the effective connectivity, than for the model where competitive interactions are turned into cooperative.

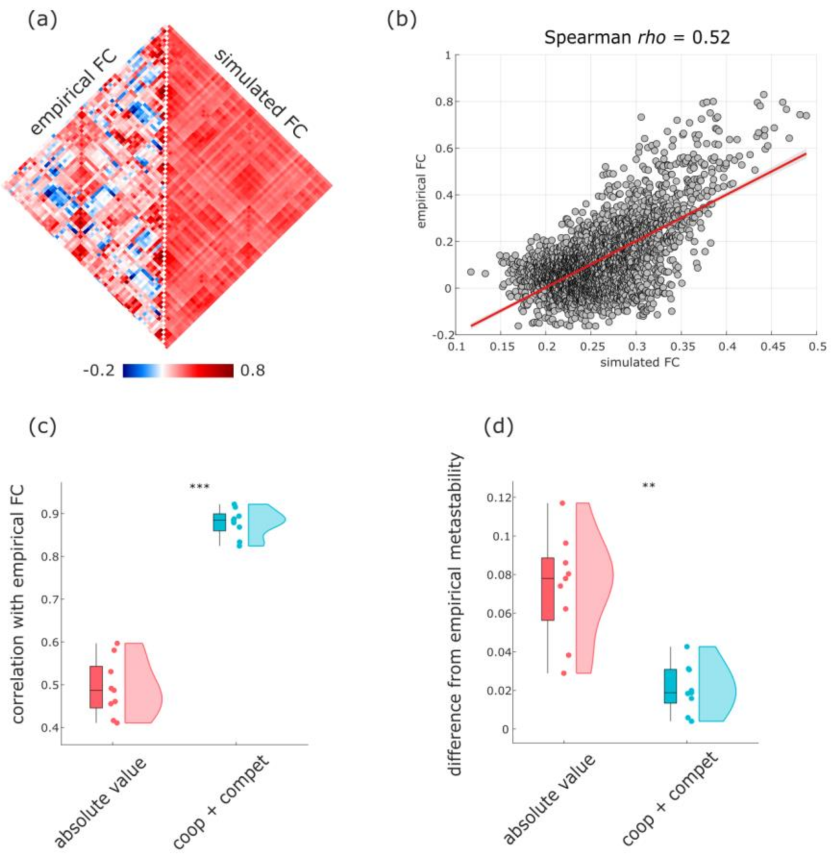


**Figure S10. Turning competitive interactions into cooperative degrades performance of mouse model.** (**a**) Group-wise empirical and simulated FC. (**b**) Correlation between group-wise empirical and simulated FC ($r = 0.52$). (**c**) Subject-wise correlations between empirical and simulated FC are significantly higher for the model including both cooperative and competitive interactions in the effective connectivity, than for the model where competitive interactions are turned into cooperative. (**d**) Subject-wise difference between empirical and simulated metastability is significantly lower (reflecting greater similarity) for the model including both cooperative and competitive interactions in the effective connectivity, than for the model where competitive interactions are turned into cooperative.

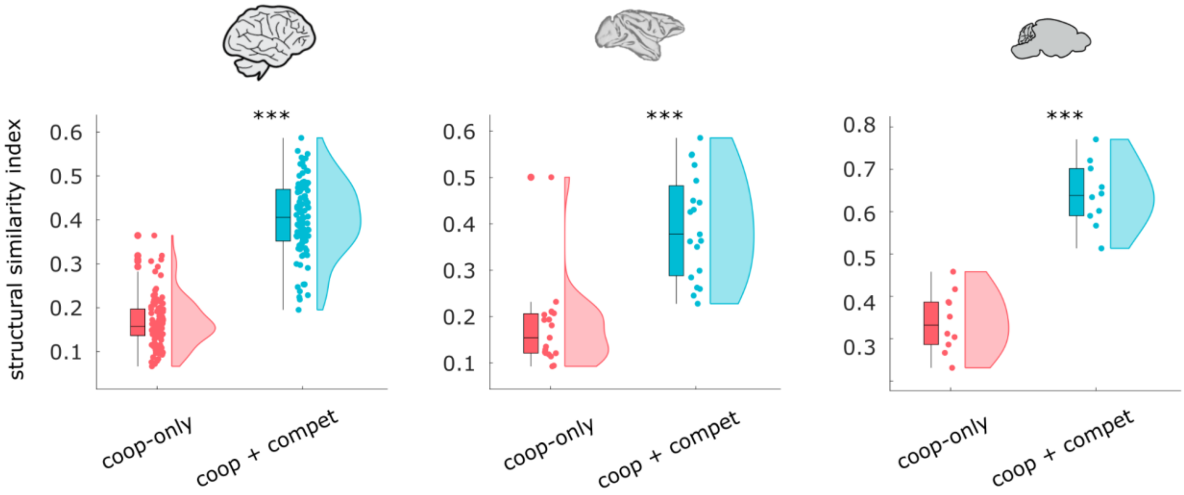


**Figure S11. Alternative fitting measure with SSIM.** In each species, the model with cooperative and competitive interactions achieves superior subject-level fit with the empirical FC, as measured by the structural similarity index (SSIM). See Supplementary Tables 1-3 for full statistical reporting.

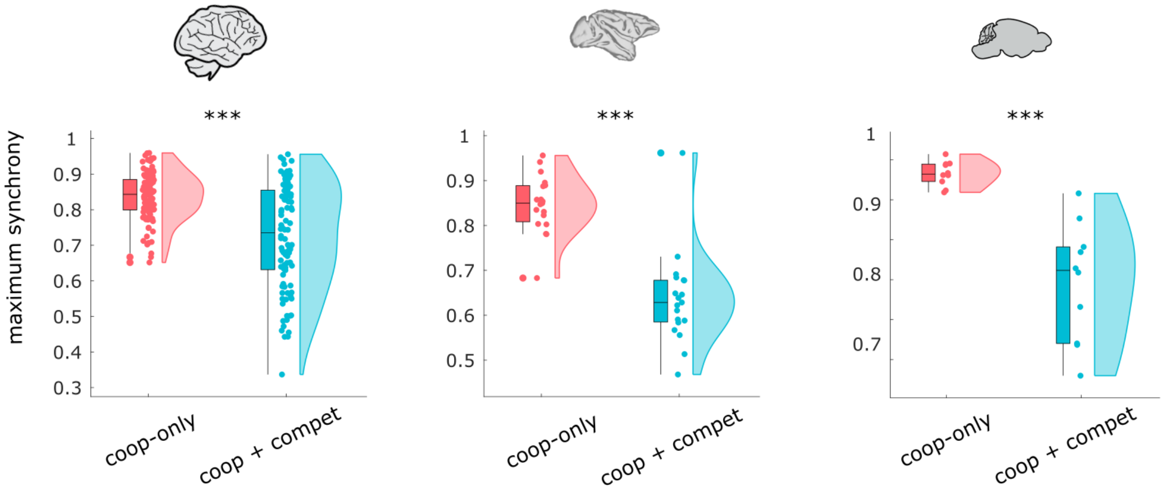


**Figure S12. Maximum synchrony.** Instantaneous synchrony is quantified by the Kuramoto order parameter of the narrowband signal; for each individual, we show its maximum observed value. The maximum synchrony is significantly higher in the model without competitive interactions. See Supplementary Tables 1-3 for full statistical reporting.

| | | | | |
|---|---|---|---|---|
| action | adaptation | addiction | anticipation | anxiety |
| arousal | association | attention | autobiographical memory | balance |
| belief | categorization | cognitive control | communication | competition |
| concept | consciousness | consolidation | context | coordination |
| decision | decision making | detection | discrimination | distraction |
| eating | efficiency | effort | emotion | emotion regulation |
| empathy | encoding | episodic memory | expectancy | expertise |
| extinction | face recognition | facial expression | familiarity | fear |
| fixation | focus | gaze | goal | hyperactivity |
| imagery | impulsivity | induction | inference | inhibition |
| insight | integration | intelligence | intention | interference |
| judgment | knowledge | language | language comprehension | learning |
| listening | localization | loss | maintenance | manipulation |
| meaning | memory | memory retrieval | mental imagery | monitoring |
| mood | morphology | motor control | movement | multisensory |
| naming | navigation | object recognition | pain | perception |
| planning | priming | psychosis | reading | reasoning |
| recall | recognition | rehearsal | reinforcement learning | response inhibition |
| response selection | retention | retrieval | reward anticipation | rhythm |
| risk | rule | salience | search | selective attention |
| semantic memory | sentence comprehension | skill | sleep | social cognition |
| spatial attention | speech perception | speech production | strategy | strength |
| stress | sustained attention | task difficulty | thought | uncertainty |
| updating | utility | valence | verbal fluency | visual attention |
| visual perception | word recognition | working memory | | |

**Figure S13. NeuroSynth terms. |** Terms that overlapped between the NeuroSynth database [115] and the Cognitive Atlas [114] were included in the analysis.
